# Comparison of the somatic TADs and lampbrush chromomere-loop complexes in transcriptionally active prophase I oocytes

**DOI:** 10.1101/2021.11.30.470320

**Authors:** Tatiana Kulikova, Antonina Maslova, Polina Starshova, Sebastian Juan Rodriguez, Alla Krasikova

## Abstract

In diplotene oocyte nuclei of all vertebrate species, except mammals, chromosomes lack interchromosomal contacts and chromatin is linearly compartmentalized into distinct chromomere-loop complexes forming lampbrush chromosomes. However, the mechanisms underlying the formation of chromomere-loop complexes remain unexplored. Here we aimed to juxtapose somatic topologically associating domains (TADs), recently identified in chicken embryonic fibroblasts, with chromomere-loop complexes in lampbrush meiotic chromosomes. By measuring 3D-distances and colocalization between linear equidistantly located genomic loci, positioned within one TAD or separated by a TAD border, we confirmed the presence of predicted TADs in chicken embryonic fibroblast nuclei. Using three-colored FISH with BAC probes we mapped equidistant genomic regions included in several sequential somatic TADs on isolated chicken lampbrush chromosomes. Eight genomic regions, each comprising two or three somatic TADs, were mapped to non-overlapping neighboring lampbrush chromatin domains – lateral loops, chromomeres or chromomere-loop complexes. Genomic loci from the neighboring somatic TADs could localize in one lampbrush chromomere-loop complex, while genomic loci belonging to the same somatic TAD could be localized in neighboring lampbrush chromomere-loop domains. In addition, FISH-mapping of BAC probes to the nascent transcripts on the lateral loops indicates transcription of at least 17 protein-coding genes and 2 non-coding RNA genes during the lampbrush stage of chicken oogenesis, including genes involved in oocyte maturation and early embryo development.

## Introduction

Spatial genome organization in various cell types is a fundamental issue in cell biology. In the cell nucleus genome is organized in hierarchical domains ranging from chromatin loops to chromosome territories (Bickmore and van Steensel 2013; Gibcus and Dekker 2013; Misteli 2020; Mirny and Solovei 2021). However, how chromatin domains identified by chromatin conformation capture, such as topologically associating domains (TADs) and A/B compartments, correlate with chromatin domains revealed by light and electron microscopy in vertebrates is still a matter of debate (Bintu *et al*. 2018; Szabo *et al*. 2020; Cremer *et al*. 2020; Trzaskoma *et al*. 2020).

TADs represent chromatin domains with higher frequency of genomic contacts, where allowed interactions between regulatory elements take place; TAD borders insulate contacts between neighboring chromatin domains (Bouwman and de Laat 2015; Razin *et al*. 2016; Dixon *et al*. 2016). From the beginning of studies, it was suggested that TADs, being several hundred kilobases in size in vertebrates, could represent structural units of interphase chromosomes. Evidence in favor of this hypothesis was obtained during *Drosophila* polytene chromosomes examination (Eagen *et al*. 2015; Ulianov *et al*. 2016). Comparison of TADs and banding pattern in polytene chromosomes demonstrated that inactive TADs mainly correspond to black bands of polytene chromosomes, while transcriptionally active regions within interTADs or active TADs, depending on resolution, coincide with interband sections (Eagen *et al*. 2015; Ulianov *et al*. 2016; Kolesnikova *et al*. 2018). It is suggested that in *Drosophila*, chromatin segregation into several types together with transcription play a crucial role in the appearance of TADs (Ulianov *et al*. 2016; Rowley *et al*. 2017).

In vertebrates, in addition to active and inactive chromatin segregation, another mechanism contributes to TAD formation. This mechanism involves generation of CTCF- and cohesin-mediated loops making the overall picture of contact chromatin domains organization more complicated (Rowley *et al*. 2017; Beagan and Phillips-Cremins 2020). In the current study, we employed giant transcriptionally active lampbrush chromosomes with a distinct chromomere-loop appearance to search for cytological equivalents of TADs in vertebrate cells.

One of the main advantages of lampbrush chromosome studies is having the opportunity to visualize transcriptionally active as well as presumably silent chromatin domains by light microscopy (Kaufmann *et al*. 2012; Macgregor 2012; Mirny and Solovei 2021). Chromomere arrays at the axes of lampbrush chromosomes represent a remarkable example of chromosome compartmentalization: gene loci tightly contact each other within individual chromomeres and are spatially separated from gene loci in neighboring chromomeres. Lampbrush chromomeres are usually defined as morphologically discrete dense chromatin domains (Vlad and Macgregor 1975). Historical aspects of lampbrush chromomeres investigation as well as recent findings can be found in a review by Herbert Macgregor (Macgregor 2012). In avian and amphibian lampbrush chromosomes, the majority of chromomeres are enriched with epigenetic markers of transcriptionally inactive chromatin (Angelier *et al*. 1986; Krasikova *et al*. 2009; Morgan *et al*. 2012; Krasikova and Kulikova 2017; Kulikova *et al*. 2020). In contrast, lateral loops emerging from lampbrush chromosome chromomeres represent sites of intense ongoing transcription (Gall and Callan 1962; Hartley and Callan 1978; Morgan 2018).

The genomic context of individual chromomeres was established only for a few examples including pericentromere and subtelomere chromomeres of amphibian and avian lampbrush chromosomes and chromomeres of chicken W chromosome that consists of tandem repeat families (Macgregor 2012). Gene and repeat content for the majority of regular lampbrush chromomeres remain unknown. Moreover, the mechanisms of the insularity of neighboring chromomeres in lampbrush chromosome axes are still unexplored.

Earlier, using mechanical microdissection followed by high-throughput sequencing we deciphered DNA sequences of a number of prominent chromomeres from chicken lampbrush macro- and microchromosomes (Zlotina *et al*. 2016, 2020). It was demonstrated that dense DAPI-positive and loop-less chromomeres could be enriched with different repetitive elements and depleted for genes while the other regular chromomeres have less uniform content. In addition, the discrepancy in organization between lampbrush chromomere-loop complexes and compact chromatin domains of somatic cells was suggested in these studies (Zlotina *et al*. 2020). However, due to uncertainty in the identification of the genomic borders of dissected chromomeres, more accurate approaches should be applied to precisely establish a correspondence between chromatin domains of interphase nuclei and lampbrush chromomere-loop complexes. FISH is widely used not only to verify Hi-C data, but to analyze spatial organization of chromatin domains in individual cells and chromosome preparations (Giorgetti and Heard 2016; Maslova and Krasikova 2021). Here, using DNA/DNA+RNA fluorescence *in situ* hybridization (FISH) we juxtaposed several earlier characterized somatic TADs with chromomere-loop complexes in chicken lampbrush chromosomes. We also confirmed the presence of selected TADs in chicken embryonic fibroblast nuclei by 3D-FISH. Our FISH data suggest that somatic TADs map to discrete lampbrush chromatin domains. Moreover, our data indicates that genes localize in the lateral loops extending from chromomeres more often than in chromomere cores. The results obtained give us a deeper insight into lampbrush chromatin domains organization.

## Material and methods

### Cell culture

Chicken embryonic fibroblasts (CEF) were collected according to the standard technique (Freshney 2010) from 10-day embryos. Cells were grown in DMEM/F12 (Biolot), supplied with 8% fetal bovine serum (Gibco), 2% chicken serum (Gibco) and 50 µg/ml gentamycine at 37°C and 5% CO2. Cells were passaged on a regular basis at 80% confluency. After the 5th passage cells were seeded on 18×18 glass coverslips (Menzel-Glaser), allowed to grow until 60-70% confluency and fixed in 4% PFA (Sigma) for 20 min.

### Preparation of lampbrush chromosomes

Lampbrush chromosomes of *Gallus gallus domesticus* (the domestic chicken) were prepared according to established protocols (Kropotova and Gaginskaya 1984; Solovei *et al*. 1992, https://projects.exeter.ac.uk/lampbrush/protocols.htm) under stereomicroscope Leica MZ16 (Leica Microsystems). Special attention was paid to preserve RNA on the samples, including labware pretreatments with RNAse inhibitors and sterilization of buffers and instruments. Preparations were centrifuged at 4000 rpm for 30 min, fixed in 2% formaldehyde in PBS for 30 min, dehydrated in ethanol series. Animals were handled according to the approval #131-04-6 from 25.03.2019 of the Ethics committee for Animal Research of St. Petersburg State University.

### FISH probes

For each analyzed region, comprising two-three consecutive TADs, three BAC-clones were chosen from the CHORI-261 chicken BAC-library (https://bacpacresources.org/chicken261.htm). The genomic distance between each BAC-clone from the triplet was as similar as possible (deviation ±17 Kb) and the linear separation of ∼500 Kb, ∼600 Kb and ∼700 Kb was tested for different regions. Within the BAC triplet at least one clone was separated from the two others by the TAD border. Details of BAC-clones used for probe generation are summarized in the **Supplementary Table 1**.

BAC DNA was isolated from *E. coli* night cultures according to the standard alkaline-lysis method. BAC DNA was labeled with dUTP-bio (DNA-Synthesis) or dUTP-dig (Roche) by nick translation (Knoll and Lichter 2005; Cremer *et al*. 2012) with DNA Polymerase I/DNase I mix (ThermoFisherScientific) or directly labeled with Atto647N nick translation labeling kit (JenaBioscience). Labeled probes were precipitated with 50 fold excess of salmon sperm DNA and dissolved in a hybridization buffer (50% formamide, 10% dextran sulfate, 2×SSC) at concentration 20 ng/μ

### 3D FISH

Permeabilization, pre-hybridization treatments, 3D-FISH and probe detection on cell preparations were performed according to previously established protocols (Cremer *et al*. 2012). Briefly, cells were permeabilized with 0.5% Triton X100 (Sigma) during 20 min at RT, incubated overnight in 20% glycerol in PBS, permeabilized by several freeze-thaw cycles in a liquid nitrogen, and finally treated with 0.1 N HCl for 5 min. Coverslips with cell monolayers were denatured in 70% formamide in 2×SSC at 70°C for 15 min, dehydrated in ice-cold ethanol series and air-dried. Probe DNA was denatured at 95°C for 10 min. Hybridization was performed at 37°C in a humid chamber for 15-40 hours. Biotin or digoxigenin-labeled probes were detected by Alexa488-conjugated streptavidin and Cy3-conjugated anti-digoxigenin antibody correspondingly. For signal amplification secondary biotinylated anti-streptavidin or Cy3-conjugated anti-mouse antibodies were applied. Finally, cells were counterstained with 1 μ l DAPI and mounted in homemade antifade solution (50 mM TrisHCl, pH=8.0, 90% glycerol, 1% DABCO).

### 2D FISH

To obtain pictures of native chromomeres, freshly prepared lampbrush chromosomes were stained with 1 μ ml DAPI in antifade solution described above and imaged with Leica DM4000 fluorescent microscope before FISH. 2D-FISH procedures were applied to lampbrush chromosome preparations according to DNA/DNA+RNA protocol preserving RNA (Solovei *et al*. 1994; Zlotina and Krasikova 2017). In particular, RNAase treatment was omitted. Chromosomes and DNA-probes were denatured separately as described for 3D-FISH. 1-5 μ drops of denatured hybridization mix were applied to the slides with denatured lampbrush chromosomes, mounted with coverslip and sealed with rubber cement. Hybridization was carried out overnight at 37°C in a humid chamber. Post-hybridization washing was performed at 60°С in 0.2×SSC. Biotin and digoxigenin were detected as described above. Slides were mounted with the antifade solution containing DAPI.

### Microscopy and image analysis

The cell preparations after 3D-FISH were analyzed by Leica TCS SP5 confocal laser scanning microscope (Leica-Microsystems) and HCX PL APO×100 objective. The microscope was adjusted and checked for the absence of chromatic shift during fluorochrome detection from different spectral regions using the TetraSpeck Fluorescent Microspheres Size Kit (ThermoFisher Scientific). Sequential scanning mode was used for each of the four channels (DAPI, Alexa488, Cy3, Atto647N) in order to prevent channel crosstalk. The voltage across the photomultiplier tubes (PMT) was set in such a way that the signals from all fluorochromes were clearly distinguishable, but there was no over-exposure in the brightest signal area (“glow over/under” software function). The dimensions of the image voxel varied in a range of 50- 80 nm in the lateral and 129-170 nm in the axial plane.

In order to increase the image contrast and to reduce noise for each of the confocal image series, signals were deconvolved in the AutoQuantX3 (Media Cybernetics) using the adaptive blind 3D- deconvolution algorithm, with 20 iterations for each fluorescence channel. To analyze 3D- distances between the centers of mass of fluorescent signals and percentage of overlapping of segmented objects Nemo software was used (Iannuccelli *et al*. 2010). During segmentation of FISH-signals in individual nuclei, the “Median filter” and the “TopHat” filters were applied, and the “Smooth Object” function was disabled. In the overwhelming majority of cases, the program made it possible to accurately segment 2 foci per nucleus (corresponding to two homologues) from each of the fluorochromes. In cases when one of the six signals per nucleus was missing due to ineffective hybridization only one of the two homologous regions was analyzed. The statistical analysis of the 3D-distance distribution and colocalization between FISH-signals was carried out using the Wilcoxon signed-rank test in the Origin 8.0.

Lampbrush chromosomes after 2D-FISH were analyzed by fluorescent microscope Leica DM 4000 (Leica Microsystems) and imaged by monochrome CCD camera (1.3 Mp resolution). Schematic drawings of hybridization patterns were made by analyzing at least 8 microphotographs for each studied chromosomal region.

## Results and Discussion

### Verification of TADs in interphase nuclei of chicken embryonic fibroblasts by 3D-FISH

TADs can be visually identified as contact-enriched squares along the main diagonal of Hi-C heatmap of a particular chromosome; however, for unbiased identification of TAD borders various mathematical algorithms are used. By implementing three different TAD calling tools (directionality index (DI), Armatus and TADTree) Fishman at al. demonstrated that in chicken embryonic fibroblasts (CEF) local contact chromatin domains revealed on a cell population-averaged Hi-C map have a nested appearance with smaller domains (subTADs) included into larger ones (TADs) (Fishman *et al*. 2019). Importantly, different TAD calling algorithms resulted in a different number and average size of domains called in the same Hi-C map, indicating the difficulty of TAD distinction by bioinformatic methods alone.

Here we used 3D-FISH on CEF with BAC-clone based probes to verify previously estimated genomic positions of TADs in several chromosomal regions. We measured 3D-distances and colocalization between linear equidistantly located loci, either positioned within one TAD or separated by a TAD border. As the contact frequency of loci interaction within a TAD is higher, the 3D distance between loci from the same TAD is expected to be shorter (Nora *et al*. 2012; Giorgetti and Heard 2016). Genomic regions containing two or three sequential TADs selected for measurement of the distance between the FISH-probes are listed in **Supplementary Table 1**. For these regions we took into account both the size of the BAC-clone DNA inserts and some TAD characteristics. Firstly, BAC-clones (average size 182 ± 36 kb, CHORI-261 BAC-library) should be linearly separated in the genome for at least twice the BAC-clone size. Linear separation of ∼500 Kb, ∼600 Kb and ∼700 Kb was tested for different genomic regions. Secondly, the verified TAD border should be called as invariant by at least two algorithms used previously. Finally, the regions containing genomic gaps or poorly assembled subregions were avoided. In total, we selected eight chromosomal regions on GGA1, GGA2, GGA4 and GGA14, which encompassed two or three neighboring TADs and were probed by different triplets of BACs (**Supplementary Table 1**).

After 3D-FISH with BAC-clone based probes for six selected genomic regions, from 14 to 32 CEF interphase nuclei (28-66 regions) were analyzed by CLSM. 3D distances between centers of mass of fluorescent foci corresponding to linearly equidistant neighboring genomic loci varied from 0.02 to 2.8 μ (**Figure 1**). 3D distances between fluorescent foci belonging to a single TAD were significantly lower than between foci belonging to neighboring TADs in three chromosomal regions – GGA1_r1, GGA4_r1 and GGA14_r1 (**Figure 1 a**′′′; **d**′′′; **f**′′′). As expected, for these regions the percent of colocalization of the foci belonging to a single TAD was higher than between the foci belonging to the neighboring TADs (**Figure 1 a^iv^**; **d^iv^**; **f^iv^**). Furthermore, we did not find significant differences in interprobe 3D distance distribution for GGA2_r2 and GGA4_r2 regions, where each BAC-probe from the triplet fell into a separate TAD (**Figure 1 c**′′′; **e**′′′). Thus our 3D-FISH data confirms the appearance of predicted TADs in individual interphase nuclei of chicken embryonic fibroblasts.

**Figure 1.**
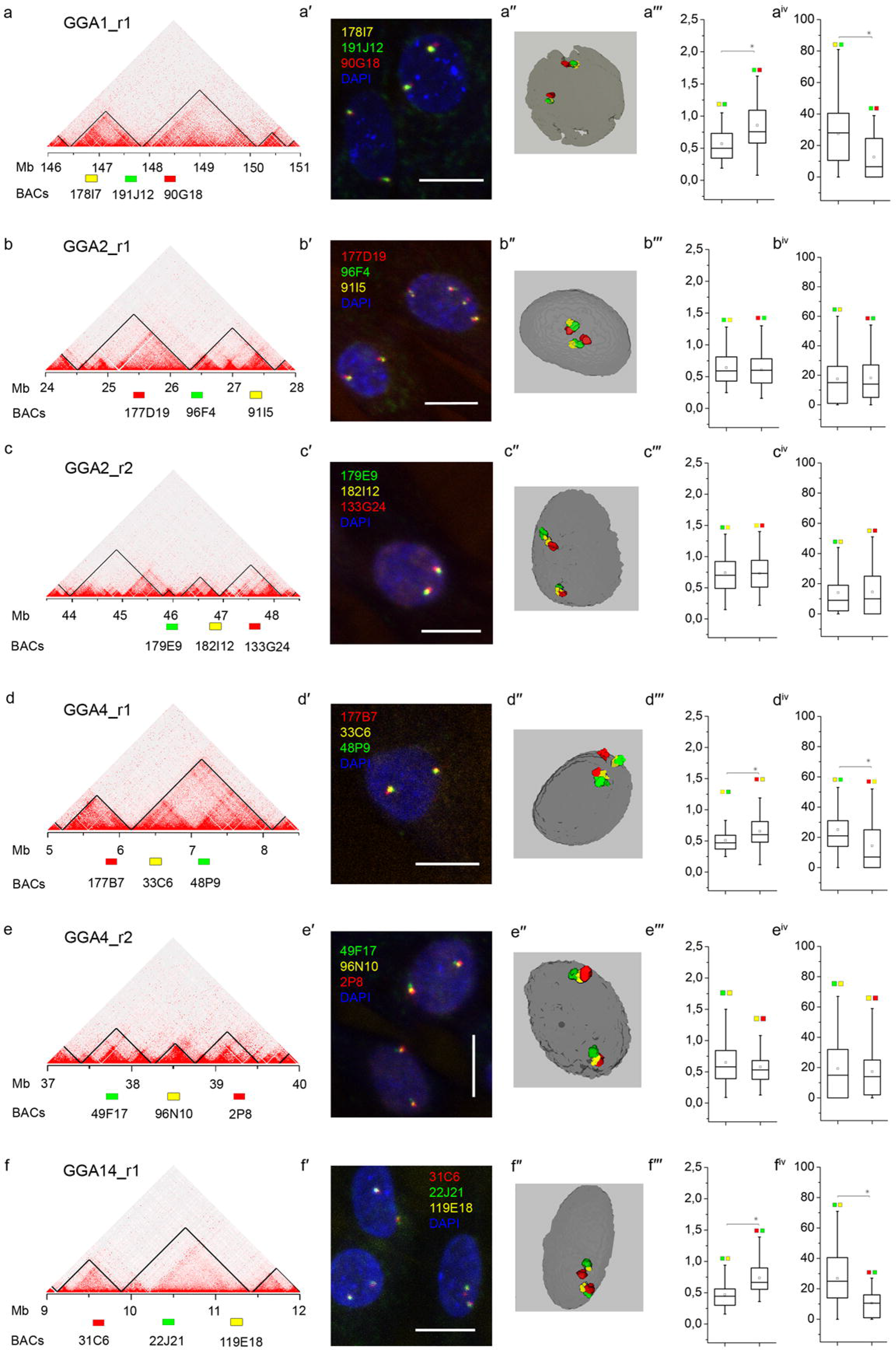
Verification of TADs in interphase nuclei of chicken embryonic fibroblasts by 3D-FISH. Measurements of 3D-distances and colocalization between linear equidistantly located genomic loci positioned within one TAD or separated by a TAD border in six chromosomal regions (GGA1_r1, GGA2_r1, GGA2_r2, GGA4_r1, GGA4_r2, and GGA14_r1). **a - f** – Hi-C heatmaps of genomic interactions within the analyzed regions in chicken embryonic fibroblasts (according to Fishman et al., 2019); positions of BAC-clones relative to TAD borders identified by directionality index algorithm are indicated. **a**′ **- f**′ – examples of CEF images after 3D-FISH with BAC-clone based probes to the analyzed regions. Nuclei are counterstained with DAPI (blue). Scale bars – 10 μm. **a**′′ **- f**′′ – 3D-surface reconstructions of FISH-signals and nuclear borders. **a**′′′ **- f**′′′ – pairwise 3D-distances between mass centers of imaged regions in a cell sample represented as boxplots. **aiv - fiv** – signal co-localization estimated by percent of signal overlap, depicted as boxplots. An asterisk denotes statistically significant differences between the probe pairs.

However, we did not find statistically significant differences in the 3D distances for the genomic loci belonging to a single or neighboring TADs in the chromosomal region GGA2_r1 (**Figure 1 b**′′′). The inconsistency between the data obtained by chromosome conformation capture, which is population-based, and loci FISH-imaging, which is single-cell based, could derive from general difference between the two methods (Dekker 2016; Giorgetti and Heard 2016). On the other hand, the discrepancy between Hi-C and 3D-FISH data for the GGA2_r1 region can be explained by inaccurate identification of TADs in this particular region by the directionality index algorithm (Dixon *et al*. 2012). Within the TAD, hybridizing with two BAC-probes in GGA2_r1, several nested domains are clearly observed (**Figure 1 b**); these nested contact domains could correspond to separate TADs. Indeed, alternative algorithms of TAD borders identification – Armatus (Filippova *et al*. 2014) and TADtree (Weinreb and Raphael 2016) – reveal 3-4 large and 4 small domains in the GGA2_r1 region (**Supplementary Figure 1**). According to these algorithms, three analyzed genomic loci lie within separate TADs which well corresponds to our 3D-FISH analysis of spatial chromatin organization in the region GGA2_r1. When interpreting 2D-FISH data within the GGA2_r1 region on lampbrush chromosome preparations, we have taken into account that in CEFs the three BAC-probes selected for this region most probably map to individual TADs.

### FISH-mapping of chicken somatic interphase TADs on lampbrush chromosomes

To juxtapose somatic interphase TADs and lampbrush chromatin domains (lateral loops and chromomeres) we mapped a set of BAC-clone based DNA-probes described above to corresponding chicken lampbrush chromosomes. Lampbrush chromosomes of the domestic chicken (*Gallus gallus domesticus*) have been characterized in detail (Ahmad 1970; Hutchison 1987; Chelysheva *et al*. 1990; Derjusheva *et al*. 2003). FISH on chicken lampbrush chromosomes enables not only to map genomic sequences with high cytogenetic resolution (Galkina *et al*. 2006), but to reveal both transcribed and untranscribed DNA sequences (Hori *et al*. 1996; Krasikova *et al*. 2006; Deryusheva *et al*. 2007; Gaginskaya *et al*. 2009; Krasikova *et al*. 2012). Here, DNA-probes to the loci belonging to the neighboring interphase TADs were mapped on lampbrush chromosomes GGA1, GGA2, GGA4 and GGA14 by 2D-FISH with preservation of RNA to reveal nascent transcripts on the lateral loops. For accurate morphological analysis of chromatin domains, a native DAPI-staining pattern was imaged before chromosome denaturation. Hybridization patterns were analyzed on the base of 5-10 FISH images for each studied chromosomal region.

#### GGA1_r1

All three DNA-probes to the region GGA1_r1 hybridized to the proximal part of the large elongated chromomere at the distal part of the q-arm of GGA1 (**Figure 2**). Fluorescent foci from the DNA-probes to the loci belonging to one TAD (178I7 and 191J12) and to the neighboring TAD (90G18) often tightly adjoined each other or even colocalized (**Figure 2 a -**a**″;** Supplementary Figure 2 a-a ). Thus within the region GGA1_r1, loci belonging to two neighboring somatic TADs were mapped to a single lampbrush chromatin domain – one elongated chromomere. In the chicken genome assembly galGal5, region GGA1_r1 is in close proximity to the previously microdissected and sequenced chromomere #16-16 (Zlotina *et al*. 2016). This chromomere belongs to a gene-poor and repeat-rich type of chromomeres (Zlotina *et al*. 2016) and combines epigenetic markers of both transcriptionally active and silent chromatin (Kulikova *et al*. 2020).

**Figure 2.**
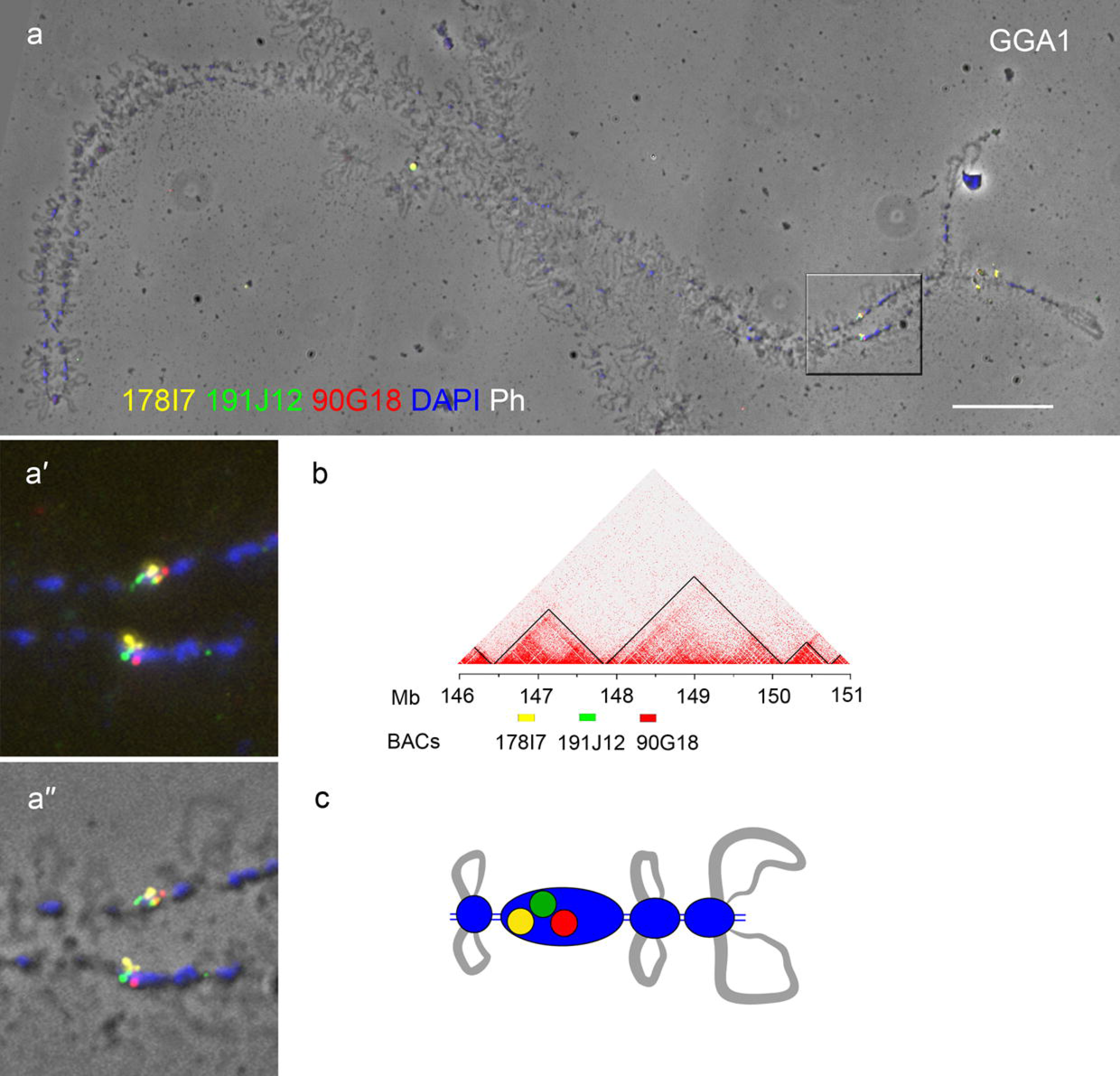
FISH-mapping of DNA-probes to the loci within two neighboring interphase TADs on chicken lampbrush chromosome 1. **a - a**′′ – FISH with BAC-clone DNA-probes to the loci belonging to two neighboring interphase TADs in the region GGA1_r1 on chicken lampbrush chromosome 1 (GGA1); **a** – lampbrush chromosome 1, fluorescent images merged with phase contrast image; **a**′, **a**′′ – enlarged region GGA1_r1, **a**′ – fluorescent image; **a**′′ – fluorescent image merged with phase contrast image. 178I7-Atto647N – yellow, 191J12-bio – green, 90G18-dig – red, DAPI – blue; scale bars: **a** – 20 μm; **a**′, **a**′′ – 10 μm. **b** – Hi-C heatmap for chicken embryonic fibroblasts (CEF) for the region GGA1_r1 (according to Fishman *et al*. 2019), positions of BAC-clones indicated with boxes of the colors corresponding to that on panels **a - a**′′. **c** – schematic drawing of hybridization pattern with DNA-probes to interphase TADs loci in the region of GGA1_r1 of lampbrush chromosome 1.

#### GGA2_r1

Genomic loci belonging to two neighboring TADs or three subTADs in the region GGA2_r1 were mapped to three separate lampbrush chromatin domains on the p-arm of GGA2 (**Figure 3**). DNA-probe 177D19 hybridized to a middle-sized chromomere. DNA-probe 96F4 hybridized with the nascent RNA transcripts within the relatively thin RNP-matrix on the pair of lateral loops extending from the next chromomere. It is worth noting that a probe to a locus hitting a transcription unit hybridizes both with the DNA of the target locus on the loop axis and with nascent RNA transcripts along the whole transcription unit downstream the locus, thus the hybridization signal can be seen along a large portion or even the whole contour of the lateral loop. DNA-probe 91I5 hybridized to the next chromomere; on some preparations, very short lateral loops extending from this chromomere also hybridized with this probe (**Figure 3 a - a**′; **Supplementary Figure 2 b - b^iv^**).

**Figure 3.**
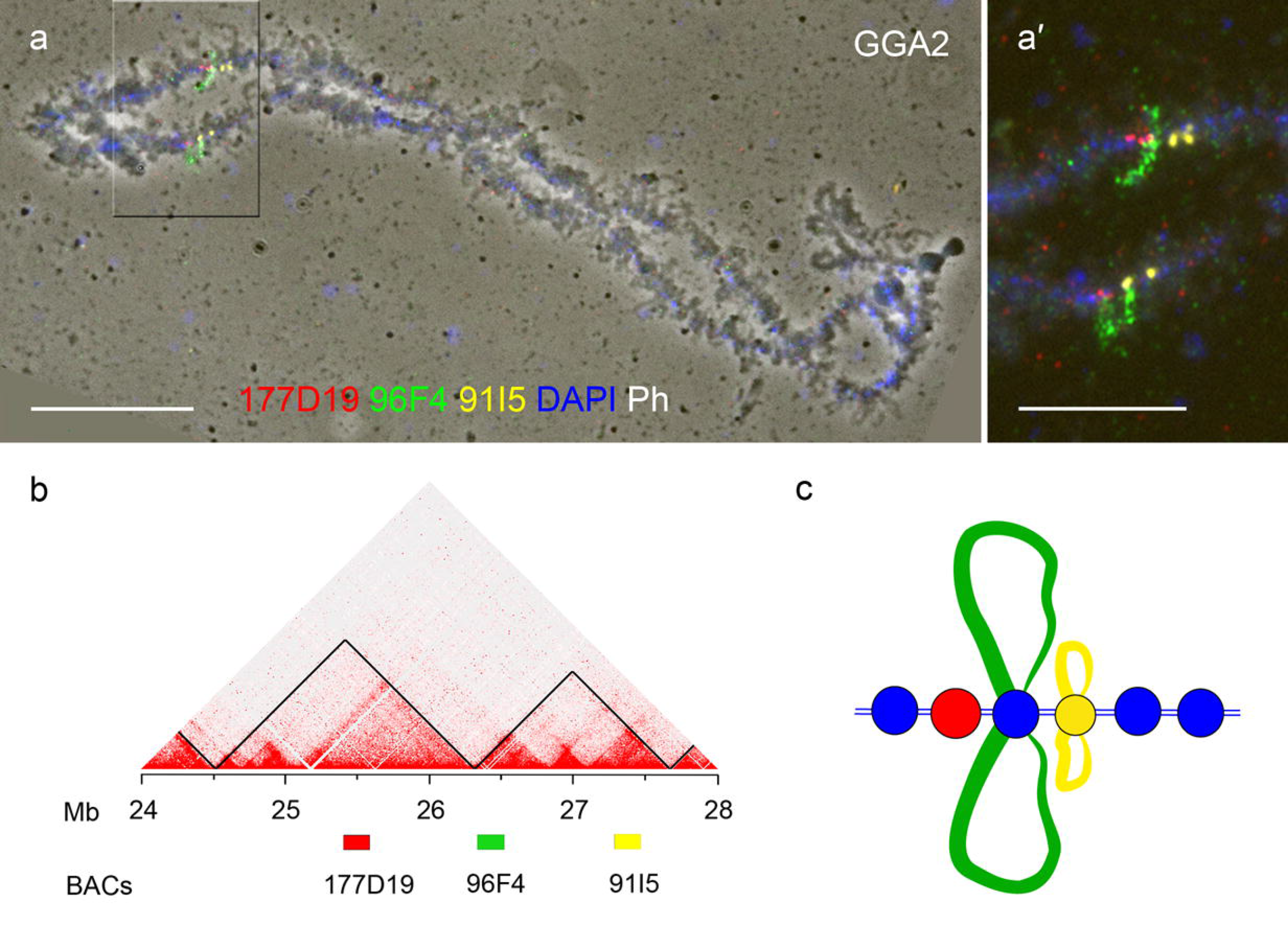
FISH-mapping of DNA-probes to the loci within two neighboring interphase TADs (three subTADs) on chicken lampbrush chromosome 2 (region GGA2_r1). **a - a**′′ – FISH with BAC-clone DNA-probes to the loci belonging to two neighboring interphase TADs (or three subTADs) in the region GGA2_r1 on chicken lampbrush chromosome 2 (GGA2); **a** – lampbrush chromosome 2, fluorescent images merged with phase contrast image; **a**′ – enlarged fluorescent image of the region GGA2_r1. 177D19-dig – red, 96F4-bio – green, 91I15-Atto647N – yellow, DAPI – blue; scale bars: **a** – 20 μm, **a**′ – 10 μm. **b** – Hi-C heatmap for chicken embryonic fibroblasts (CEF) for the region GGA2_r1 (according to Fishman *et al*. 2019), positions of BAC-clones indicated with boxes of the colors corresponding to that on panels **a - a**′. **c** – schematic drawing of hybridization pattern with DNA-probes to interphase TADs loci in the region of GGA2_r1 of lampbrush chromosome 2.

#### GGA2_r2

Region GGA2_r2 mapped to a cluster of dense pericentromeric chromomeres bearing very short lateral loops on the p-arm of GGA2. Similar to GGA1_r1, each of three DNA- probes to the loci belonging to three sequential TADs within GGA2_r1 hybridized to three sequential chromomeres (**Figure 4**). DNA-probes 179E9-bio and 182I12 hybridized exclusively to a single chromomere or to two neighboring chromomeres divided by the interchromomeric axis. DNA-probe 133G24 hybridized to the next one or two chromomeres and also to a pair of short lateral loops extending from the distal chromomere (**Figure 4 a**, **a**′′; **Supplementary Figure 2 c - c^iv^**).

**Figure 4.**
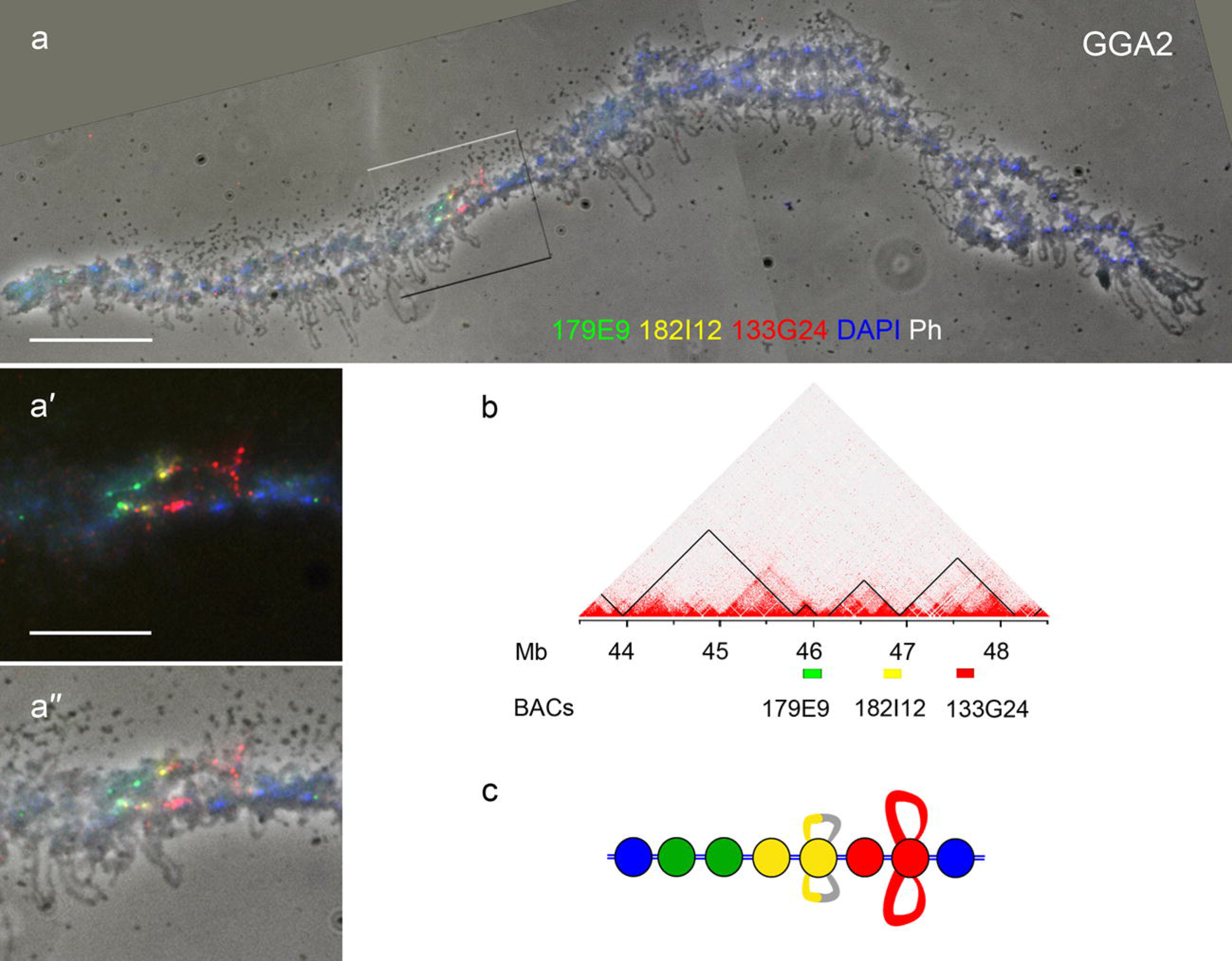
FISH-mapping of DNA-probes to the loci within three neighboring interphase TADs on chicken lampbrush chromosome 2 (region GGA2_r2). **a - a**′′ – FISH with BAC-clone DNA-probes to the loci belonging to three neighboring interphase TADs in the region GGA2_r2 on chicken lampbrush chromosome 2 (GGA2); **a** – lampbrush chromosome 2, fluorescent images merged with phase contrast image; **a**′, **a**′′ – enlarged fluorescent (**a**′) and fluorescent image merged with phase contrast image (**a**′′) of the region GGA2_r2. 179E9-bio – green, 182I12-Atto647N – yellow, 133G24-dig – red, DAPI – blue; scale bars: **a** – 20 μm, **a**′, **a**′′ – 10 μm. **b** – Hi-C heatmap for chicken embryonic fibroblasts (CEF) for the region GGA2_r2 (according to Fishman *et al*. 2019), positions of BAC-clones indicated with boxes of the colors corresponding to that on panels **a - a**′′. **c** – schematic drawing of hybridization pattern with DNA-probes to interphase TADs loci in the region of GGA2_r2 of lampbrush chromosome 2.

#### GGA4_r1

Region GGA4_r1 was mapped to the terminus of GGA4 p-arm – a region of tiny chromomeres with long lateral loops (**Figure 5**). P-arm of GGA4 evolutionary originates from one of the microchromosomes (Shibusawa *et al*. 2004; Griffin *et al*. 2007), which generally demonstrate higher gene density than macrochromosomes (Burt 2002). All three DNA-probes to the loci of two neighboring TADs fell into a single chromomere-loop complex. DNA-probe 177B7 hybridized with thick RNP-matrix along a transcription unit encompassing a large part of the axes of a pair of lateral loops emerging from a chromomere, which hybridized with the next DNA-probe — 33C6. Locus of the right subTAD (DNA-probe 48P9) was mapped to the dot-like chromatin nodule forming between two transcription units on the axes of a pair of long lateral loops emerging from the same chromomere (**Figure 5 a - a^iv^**; **Supplementary Figure 2 d - d^iv^**). Such transcriptionally silent chromatin structures on the loop axes correspond to earlier characterized untranscribed spacers between neighboring transcription units (Angelier *et al*. 1986; Morgan *et al*. 2012).

**Figure 5.**
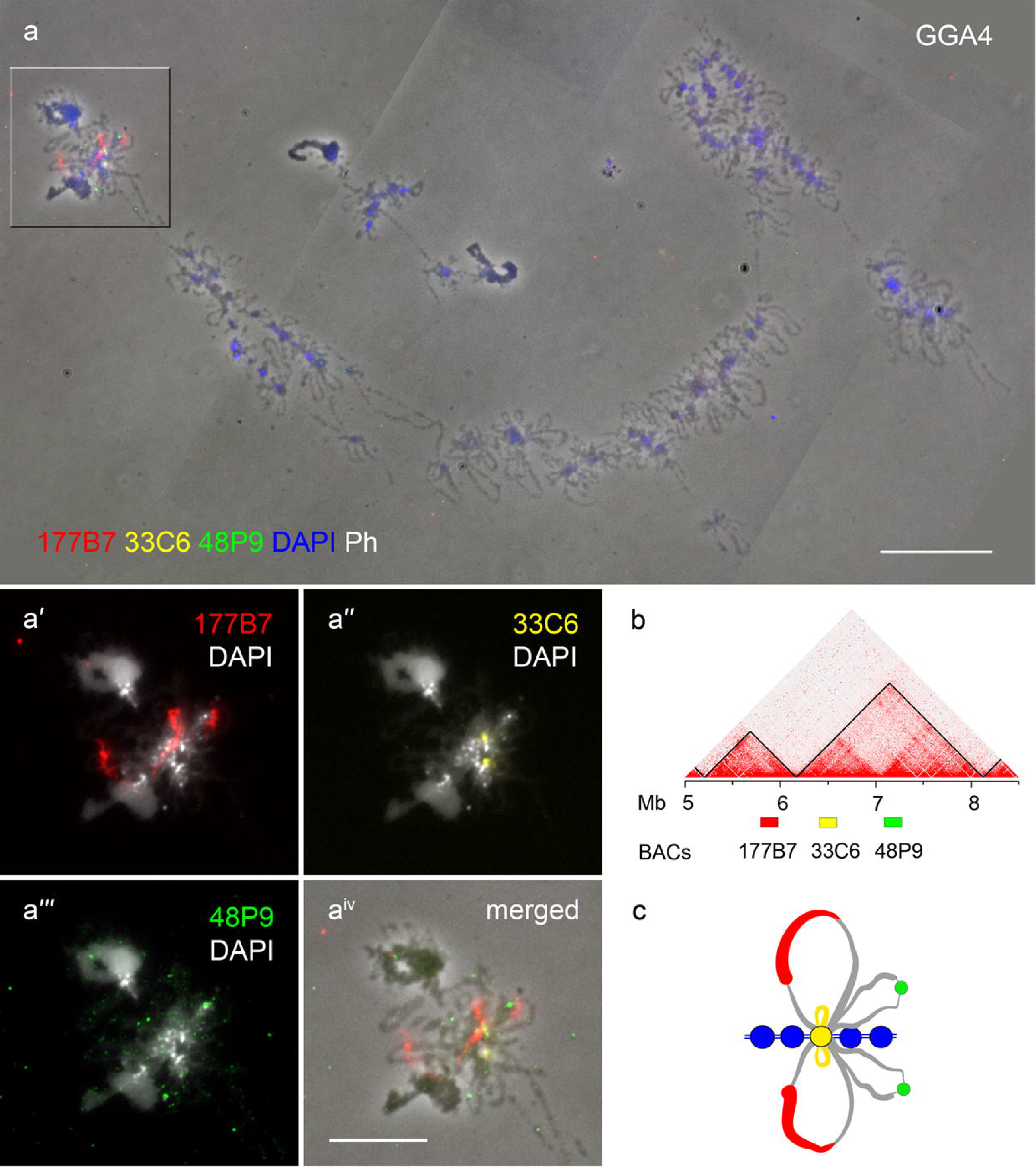
FISH-mapping of DNA-probes to the loci within two neighboring interphase TADs on chicken lampbrush chromosome 4 (region GGA4_r1). **a - aiv** – FISH with BAC-clone DNA-probes to the loci belonging to two neighboring interphase TADs in the region GGA4_r1 on chicken lampbrush chromosome 4 (GGA4); **a** – lampbrush chromosome 4, fluorescent images merged with phase contrast image, 177B7-dig – red, 33C6- Atto647N – yellow, 48P9-bio – green, DAPI – blue, scale bar – 20 μm; **a**′ **- aiv** – enlarged images of the region GGA4_r1, DNA-probes colored as on **a**, DAPI – white, **aiv** – fluorescent images merged with the phase contrast image, DAPI – blue, scale bar – 10 μm. **b** – Hi-C heatmap for chicken embryonic fibroblasts (CEF) for the region GGA4_r1 (according to Fishman *et al*. 2019), positions of BAC-clones indicated with boxes of the colors corresponding to that on panels **a - aiv**. **c** – schematic drawing of hybridization pattern with DNA-probes to interphase TADs loci in the region of GGA4_r1 of lampbrush chromosome 4.

#### GGA4_r2

Region GGA4_r2 from the middle of GGA4 q-arm mapped to the region of “weak” chromomeres presumably tending to lose longitudinal condensin molecular clips, which results in chromomere splitting into two or more segments and formation of so-called “double-loop bridges” (Chelysheva *et al*. 1990; Gaginskaya *et al*. 2009; Macgregor 2012). DNA-probes to the loci of three sequential TADs hybridized to three sequential lampbrush chromatin domains. DNA-probe 49F17 hybridized to the RNP-matrix of long lateral loops, whereas DNA-probes 96N10 and 2P8 hybridized with both chromatin of small chromomeres and RNP-matrix of lateral loops emerging from these chromomeres (**Figure 6**). An example of “double loop bridge” structures formed by chromomere-loop complexes at the region GGA4_r2 is presented in Supplementary Figure 3.

**Figure 6.**
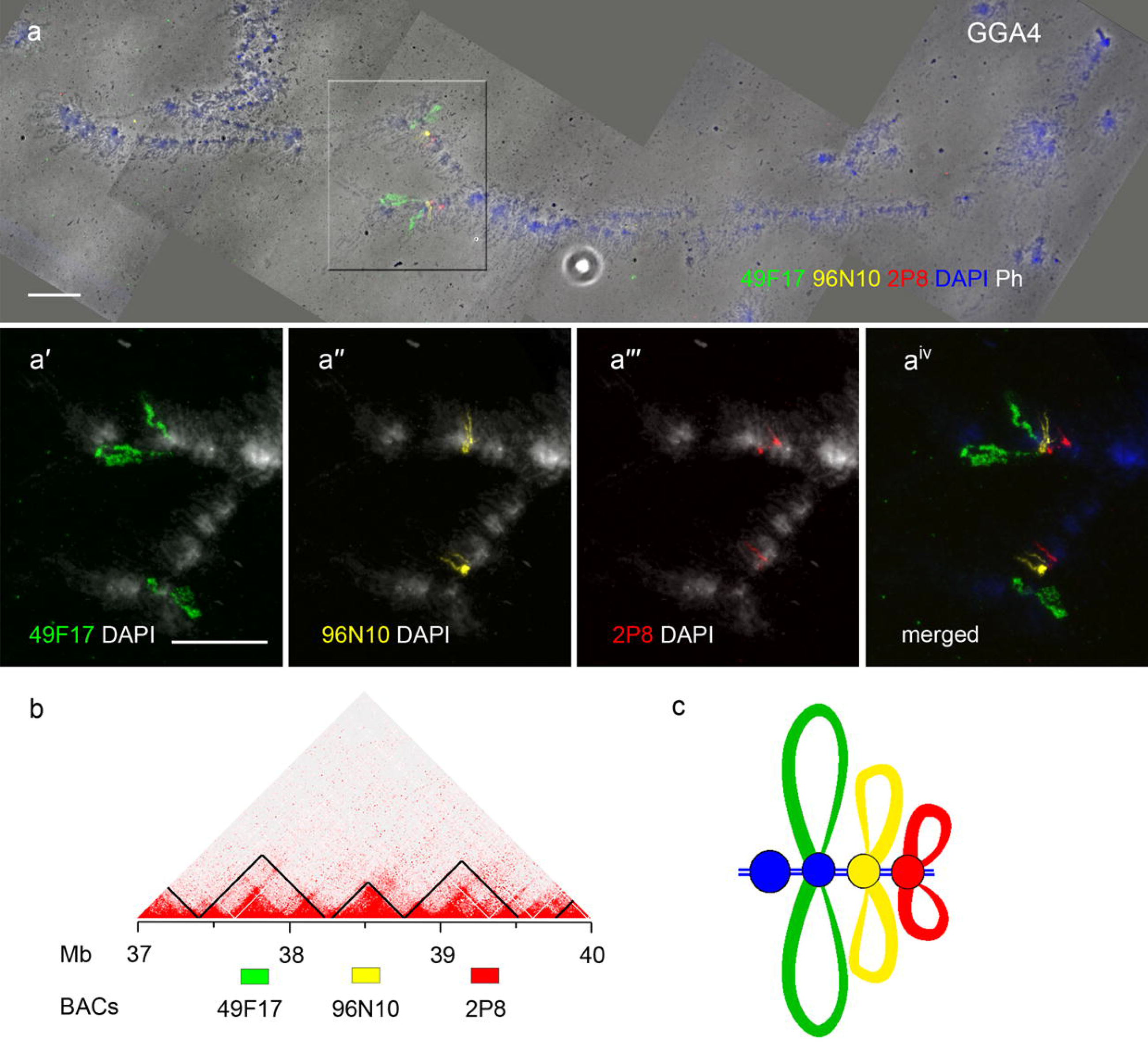
FISH-mapping of DNA-probes to the loci within three neighboring interphase TADs on chicken lampbrush chromosome 4 (region GGA4_r2). **a - aiv** – FISH with BAC-clone DNA-probes to the loci belonging to three neighboring interphase TADs in the region GGA4_r2 on chicken lampbrush chromosome 4 (GGA4); **a** – lampbrush chromosome 4, fluorescent images merged with phase contrast image, 49F17-bio – green, 96N10-Atto647N – yellow, 2P8-dig – red, DAPI – blue, scale bar – 20 μm; **a**′ **- aiv** – enlarged images of the region GGA4_r2, DNA-probes colored as on **a** separately merged with DAPI (white) (**a**′ **- a**′′′), **aiv** – merged image (DAPI – blue), scale bar – 10 μm. **b** – Hi-C heatmap for chicken embryonic fibroblasts (CEF) for the region GGA4_r2 (according to Fishman *et al*. 2019), positions of BAC-clones indicated with boxes of the colors corresponding to that on panels **a - aiv**. **c** – schematic drawing of hybridization pattern with DNA-probes to interphase TADs loci in the region of GGA4_r2 of lampbrush chromosome 4.

#### GGA4_r3

Region GGA4_r3 is located 800 kb distally from the region GGA4_r2 and is also characterized by long lateral loops, small chromomeres and frequent formation of double loop bridges. Interestingly, DNA-probes 108G8 and 34P18 to the loci belonging to one TAD hybridized with RNP-matrix of lateral loops apparently emerging from one chromomere. DNA- probe 47F11 to the loci of the neighboring TAD hybridized with both transcription units on a pair of small lateral loops and with chromatin nodules on the axes of these loops emerging from the next chromomere (**Supplementary Figure 4 a - d**; **Supplementary Figure 5 a - d**). Thus, loci belonging to two neighboring TADs in the region GGA4_r3 were mapped to the parts of two neighboring chromomere-loop complexes.

#### GGA4_r4

Region GGA4_r4 was mapped to the proximal part of the region with relatively dense chromomeres and short lateral loops at the distal of GGA4 q-arm. DNA-probes to the loci belonging to one TAD apparently hybridized with the parts of two neighboring chromomere-loop complexes: DNA-probe 14D4 hybridized with a chromomere and a pair of short lateral loops emerging from it, while DNA-probe 139O9 hybridized with the next small chromomere. DNA-probe 164E15 hybridized with the RNP-matrix on the pair of lateral loops emerging from the chromomere that hybridized with the DNA-probe 139O9 (**Supplementary Figure 4 a**, **e -g**; **Supplementary Figure 5 a**, **e-g**). In the region GGA4_r4 two neighboring loci belonging to two neighboring TADs were mapped to the non-overlapping parts of one chromomere-loop complex.

#### GGA14_r1

Region GGA14_r1 of the microchromosome GGA14 was mapped to the proximal part of the q-arm with small chromomeres and long lateral loops (**Figure 7**). Three DNA-probes to the loci of two neighboring TADs hybridized to three sequential chromomere loop complexes. DNA-probe 31C6 was mapped to a chromomere-loop complex: a pair of lateral loops and a chromomere from which they emerge. DNA-probes 22J21 and 119E18 regardless their belonging to the same somatic TAD were mapped to the separate lampbrush chromatin domains: 22J21 mapped to a chromomere and a pair of lateral loop, whereas 119E18 mapped to the RNP-matrix along the transcription unit encompassing a part of the axis of a pair of very long lateral loops (**Figure 7 a - a^v^**; **Supplementary Figure 2 e - e^iv^)**. It is worth noting that previous BAC-clone mapping (Zlotina *et al*. 2012) as well as the results of the present study revealed that gene order in GGA14 in the chicken genome assembly version galGal5 is reversed relative to the centromere position.

**Figure 7.**
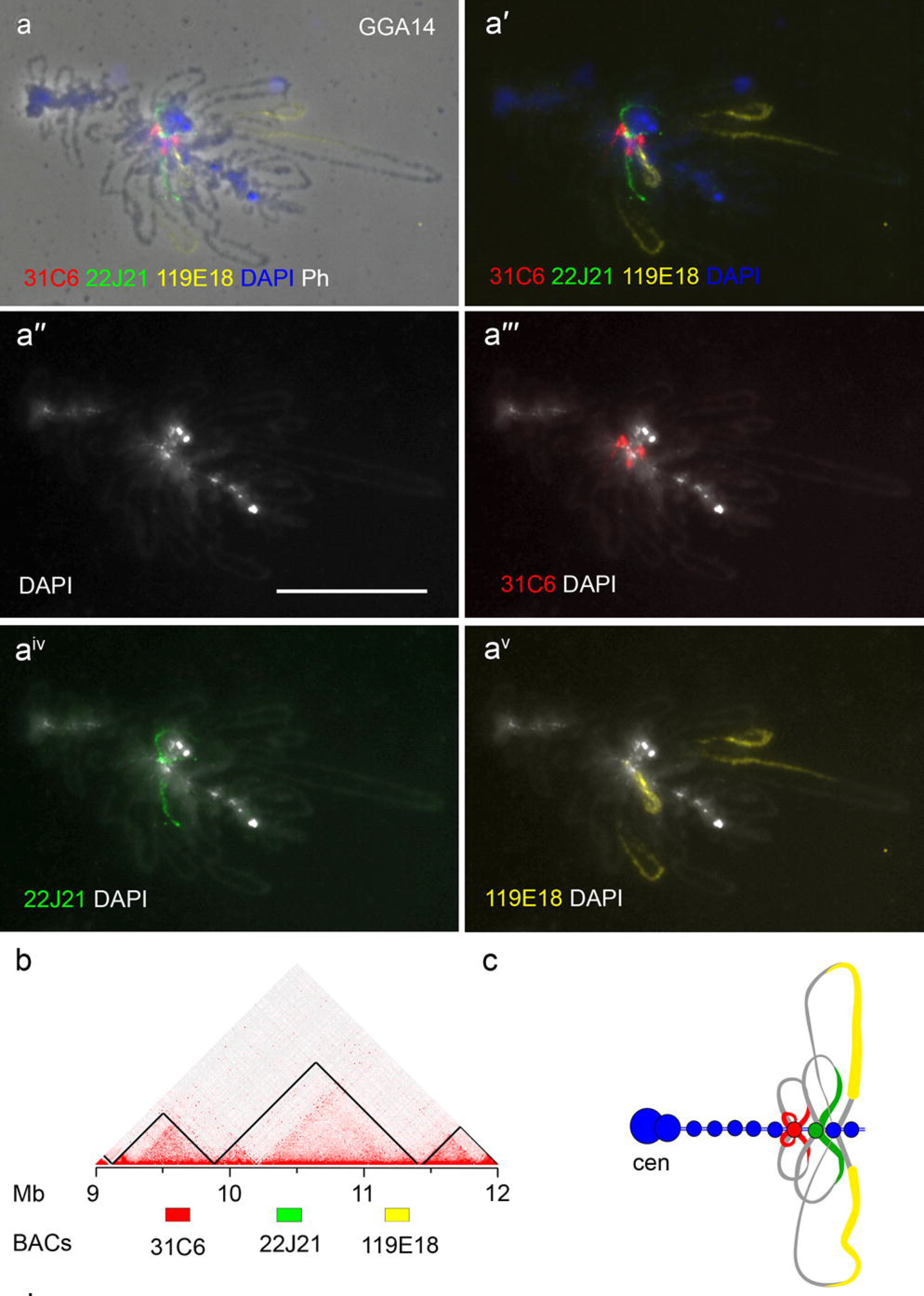
FISH-mapping of DNA-probes to the loci within two neighboring interphase TADs on chicken lampbrush chromosome 14 (region GGA14_r1). **a - av** – FISH with BAC-clone DNA-probes to the loci belonging to two neighboring interphase TADs in the region GGA14_r1 on chicken lampbrush chromosome 14 (GGA14); **a** – fluorescent images merged with phase contrast image, **a**′ – merged fluorescent images, 31C6-dig – red, 22J21-bio – green, 119E18-Atto647N – yellow, DAPI – blue, scale bar – 20 μm; **a**′′ **- av** – DNAprobes colored as on **a** separately merged with DAPI (white), scale bar – 20 μm. **b** – Hi-C heatmap for chicken embryonic fibroblasts (CEF) for the region GGA14_r1 (according to Fishman et al., 2019), positions of BAC-clones indicated with boxes of the colors corresponding to that on panels **a - av**. **c** – schematic drawing of hybridization pattern with DNA-probes to interphase TADs loci in the region of GGA14_r1 of lampbrush chromosome 14.

The obtained results enabled us to juxtapose 19 somatic interphase TADs recently identified in chicken embryonic fibroblasts with chromomere-loop complexes within eight regions of corresponding lampbrush chromosomes. We conclude that genomic loci of neighboring somatic TADs could localize in one lampbrush chromomere-loop complex or even in one chromomere, while genomic loci belonging to the same somatic TAD could be revealed in neighboring lampbrush chromomere-loop domains. Overall, the pattern of somatic TAD regions localization relative to lampbrush chromomere-loop complexes was quite diverse and depended on genomic region characteristics.

### Analysis of gene content in the sequences mapped to lampbrush chromomeres or lateral loops

FISH-mapping of BAC-clones to isolated lampbrush chromosomes enables to reveal the transcriptional status of the mapped genomic sequences (transcribed, untranscribed or partially transcribed) (Zlotina and Krasikova 2017). Ongoing transcription on the lateral loops of avian lampbrush chromosomes was demonstrated by intense and rapid RNase-sensitive incorporation over the lampbrush chromosomes and oocyte nucleoplasm of [3H]uridine after intraperitoneal injection (Callebaut 1973), as well as by incorporation into the nascent RNA of bromo-uridine triphosphate microinjected directly into the oocyte (Deryusheva *et al*. 2007; Kulikova *et al*. 2016).

Here we analyzed the gene content in the genomic loci mapped to transcriptionally active and/or inactive chromatin domains of chicken lampbrush chromosomes. 16 of 24 analyzed genomic loci mapped to the transcriptionally active chromatin domains: 7 mapped exclusively to transcription units on the lateral loops and 9 mapped to chromomere-loop complexes (both chromomere and emerging lateral loops). 8 of 24 genomic loci mapped to transcriptionally inactive chromatin domains: chromomeres or a chromatin nodule at the axes of lateral loops. In total, 24 BAC-clones selected as DNA-probes to genomic loci within the neighboring TADs overlapped with 37 annotated genes – 24 protein coding genes and 13 non-coding RNA genes (12 uncharacterized long non-coding RNAs and 1 miRNA) (data summarized in **Supplementary Table 2**). 4 of 8 BAC-clones that have been mapped to the transcriptionally inactive chromatin domains (chromomeres and a chromatin nodule) overlapped with the genomic regions lacking annotated gene sequences. BAC-clones which were mapped to chromomere-loop complexes overlapped with 17 annotated gene sequences, 8 of which were protein coding (**Figure 8**; **Supplementary Table 2**). Further analysis is needed to reveal which of these sequences are transcriptionally active. BAC-clones which were mapped exclusively to transcription units with the only one exception (34P18) overlapped with protein coding genes (**Figure 8**; **Supplementary Table 2**).

**Figure 8.**
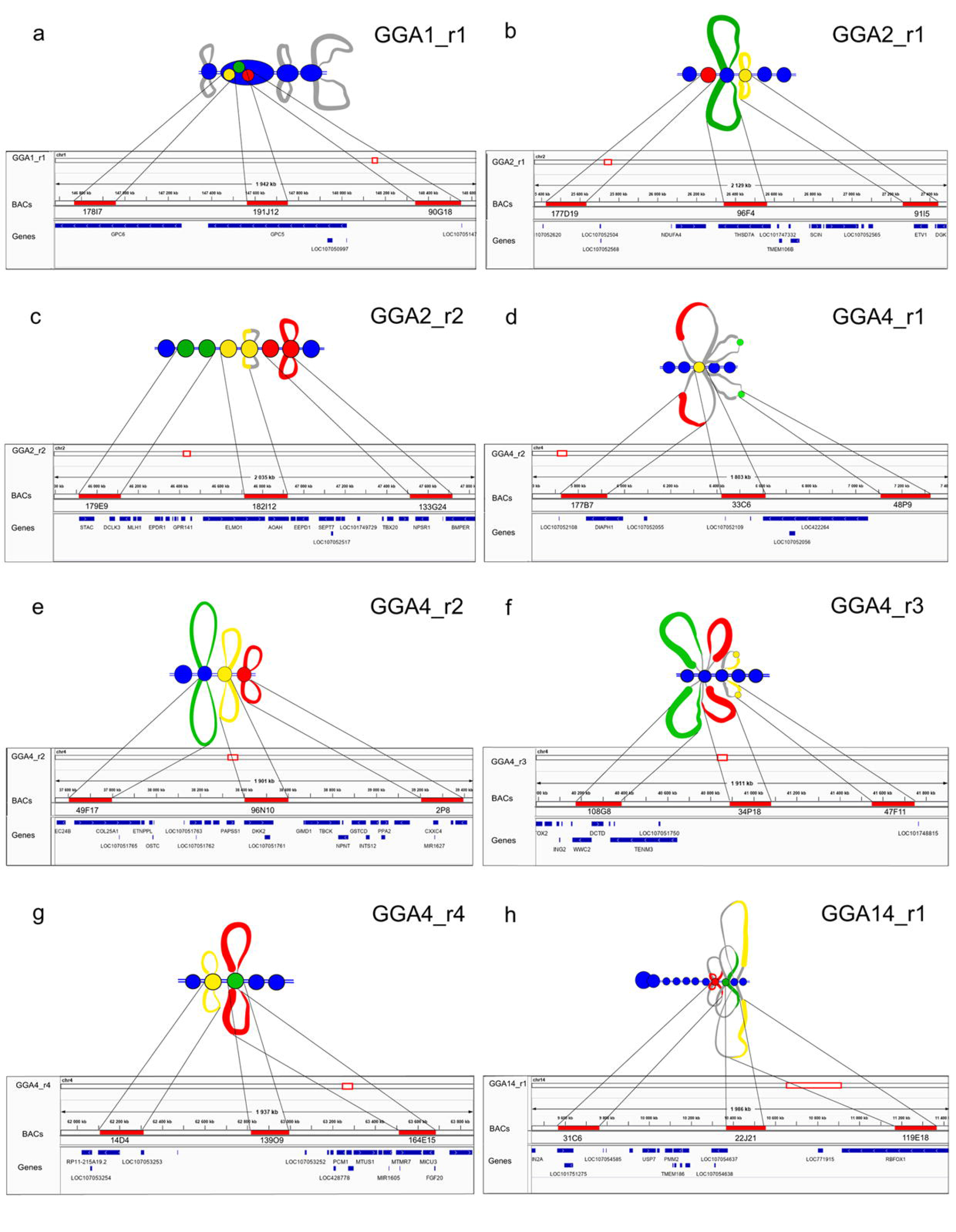
Correspondence of lampbrush chromatin domains to annotated gene sequences. **a - h** – schematic drawings of DNA-probes hybridization patterns on chicken lampbrush chromosome regions and snapshots of the Integrative Genome Viewer displaying regions occupied by BAC-clones and tracks for annotated gene sequences (chicken genome assembly version galGal5).

In total, by FISH-mapping we established transcription of 12 protein-coding genes and 2 non-coding RNA genes. BAC-clone 96F4 that was mapped to a transcription unit in the region GGA2_r1 (**Figure 3**; **Supplementary Figure 2 b - b^iv^**) completely overlaps with the gene *THSD7A* (thrombospondin, type I, domain containing 7A). BAC-clone 177B7, which was mapped to transcription units in the region GGA4_r1 (**Figure 5**; **Supplementary Figure 2 d - d^iv^**), overlaps by 60% with the gene encoding *DIAPH2* (diaphanous-related formin 2) – protein involved in ovarian development and ovarian follicle differentiation in mammals (Bione *et al*. 1998; Mandon-Pépin *et al*. 2003). In the region GGA4_r2 BAC-clone 49F17 that was mapped to the lateral loop (**Figure 6**; **Supplementary Figure 3**) overlaps with the gene *COL25A1* (collagen type XXV alpha 1 chain), which is upregulated in eggshell gland and found to be one of the key genes affecting eggshell quality in hen (Yang *et al*. 2020). BAC-clone 108G8 which was mapped to transcription units in the region GGA4_r3 overlaps with 4 genes: 2 long non-coding RNA genes – *LOC107051751* and *LOC107051752* and 2 protein-coding genes – *DCTD* (dCMP deaminase) and *WWC2* (WW and C2 domain containing 2). WWC2 was recently found to regulate cell division in both early embryo and oocyte in metaphase II of meiosis (Virnicchi *et al*. 2020). BAC-clone 34P18 did not overlap with any annotated gene sequences and could comprise new genes, most probably non-coding RNA genes. BAC-clone 164E15 in the region GGA4_r4 overlapped with 6 protein coding genes: *MTMR7* (myotubularin related protein 7); *VPS37A* (VPS37A subunit of ESCRT-I); *CNOT7* (CCR4-NOT transcription complex subunit 7); *ZDHHC2* (zinc finger DHHC-type containing 2); *MICU3* (mitochondrial calcium uptake family member 3) and *FGF20* (fibroblast growth factor 20). Phosphatidylinositol phosphatase MTMR7 is expressed in Shan Ma ducks ovary (Zhu *et al*. 2016). *CNOT7* mRNA is a maternal RNA which translates to a key component of deadenylation machinery involved in maternal mRNA clearance (Ma *et al*. 2015; Mishima and Tomari 2017). Ortholog of *FGF20* in Nile tilapia specifically expresses in female gonads and its mRNA accumulates in the cytoplasm of growing oocytes (Sun *et al*. 2012). BAC-clone 119E18 which was mapped to the long transcription unit in the region GGA14_1 (**Figure 7**) completely overlaps with the large (746.054 kb) protein-coding gene *RBFOX1* (RNA binding fox-1 homolog 1). Nuclear isoform of RBFOX1 protein regulates alternative splicing, whereas the cytoplasmic isoform regulates mRNA stability and is involved in oocyte commitment and meiotic entry in Drosophila and brain development in humans (Carreira-Rosario *et al*. 2016; Conboy 2017).

Additionally, we considered 6 BAC-clones that were used in the studies of chicken centromeres and were mapped unequivocally to the transcription units on the lateral loops of chicken lampbrush chromosomes 1, 3, 11, 12 and 14 (Zlotina *et al*. 2012). Mapping two of these BAC-clones to the nascent RNA on the lateral loops was additionally verified by RNA-FISH (Krasikova *et al*. 2012). We analyzed the positions of the genetic markers of these BAC-clones relative to the annotated gene sequences in chicken genome assembly version galGal5. Five of 6 genetic markers fell into the protein coding genes indicating their expression in the oocyte nucleus (data summarized in **Supplementary Table 3**). One of the genes mapped to a transcription unit is *PDE3A* (phosphodiesterase 3A) encoding an essential factor of meiotic resumption in mammalian oocytes (Mehlmann 2013). In total, our analysis revealed 17 protein coding genes overlapping with BAC-clones which hybridize with nascent transcripts on the lateral loops of chicken lampbrush chromosomes.

## Conclusions

Direct visualization of chromatin domains combined with FISH enabled us to compare several somatic TADs with chromomere-loop complexes of meiotic transcriptionally active lampbrush chromosomes. Our data indicates that somatic TAD loci correspond to discrete lampbrush chromomere and lateral loop domains. With exclusion of the GGA1_r1 region where two interphase TADs map to a single chromomere, in seven other studied regions interphase TADs or subTADs map to non-overlapping lampbrush chromatin domains (either chromomere, or lateral loop, or chromomere-loop domain). Earlier mechanical microdissection of chromomeres in chicken lampbrush chromosome 4 followed by high-throughput sequencing of dissected DNA suggested that individual lampbrush chromomere-loop complexes could combine several DI TADs (on average 2.5 DI TADs per chromomere-loop complex) (Zlotina *et al*. 2020). According to FISH mapping of dissected regions, probes to chromomere-loop complexes obtained by mechanical microdissection combine both transcriptionally active and inactive chromatin domains.

A functional link between three-dimensional genome organization and gene expression has been established (Goetze *et al*. 2007; Holwerda and Laat 2012; Buchwalter *et al*. 2019). One TAD could comprise one, several, or many genes. TADs could be responsible for coordinated regulation of encompassed genes (Lupiáñez *et al*. 2015; Symmons *et al*. 2016). Here we describe two examples where genes from one somatic TAD are transcribed during the lampbrush stage while being localized in different lateral loop pairs (regions GGA4_r3 and GGA14_r1). In another example, genes from one somatic TAD are differentially expressed during the lampbrush stage of oogenesis being localized either in lateral loops or in a chromomere (region GGA4_r4). Interestingly, two genomic loci – 96F4 in the GGA2_r1 region and 119E18 in the GGA14_r1 region –, localizing close to the somatic TAD border, map to transcription units on the lateral loops of lampbrush chromosomes.

The results obtained also suggest higher gene content in transcriptionally active chromatin domains (lateral loops) of chicken lampbrush chromosomes. FISH-mapping of BAC probes to the nascent transcripts on the lateral loops indicates transcription of at least 17 protein-coding genes and 2 non-coding RNA genes during the lampbrush stage of chicken oogenesis. These transcripts could be used for oocyte maturation and/or early embryo development according to data on maternal RNA accumulation during the lampbrush stage of avian oogenesis (Malewska and Olszań ka 1999). We also assume that at least 8 protein-coding genes essential for oocyte maturation, ovarian development and early stages of embryogenesis are differentially expressed during the lampbrush stage of chicken oogenesis.

Despite overall similarity of TADs between different tissues and cell lines, local contact chromatin domains could be partially restructured during germ line development and oogenesis. We believe that more precise chromosome conformation capture approaches would help to reveal basic characteristics of chromomere organization in lampbrush chromosomes as well as mechanisms of their insulation. FISH mapping of TADs identified by Hi-C of individual nuclei at the lampbrush stage of oogenesis will enable to establish a correspondence between local contact-enriched self-interacting chromatin domains and their cytological equivalents in vertebrates. Furthermore, lampbrush chromosomes represent a promising model for comprehensive studies of chromatin domains in the context of their transcriptional activity with high cytological resolution.

## Acknowledgments

The research was supported by the Russian Science Foundation (grant #19-74-20075) and was performed using the equipment of the Resource Center “Molecular and Cell Technologies” (Saint-Petersburg State University).

## Author Contributions

AK conceived the study and supervised the project. AM performed 3D-FISH on chicken embryonic fibroblasts, image analysis and 3D-measurments. TK, PS and SR performed 2D-FISH on isolated lampbrush chromosomes. TK, AK and SR performed bioinformatic analysis of mapped genomic sequences. TK, AM and AK drafted the manuscript. All authors have read and agreed to the final version of the manuscript.

## Supplementary Materials I

**Supplementary Table 1**. Positions of BAC-clones within studied regions in the chicken genome assembly version galGal5.

**Supplementary Table 2.** Summary of FISH-mapping data of BAC-clone DNA-probes to lampbrush chromatin domains with information on gene content according to chicken genome assembly version galGal5.

**Supplementary Table 3.** Summary of FISH-mapping data of BAC-clone DNA-probes to lampbrush lateral loops obtained from previous studies (Krasikova *et al.* 2012; Zlotina *et al.* 2012) with information on gene content according to chicken genome assembly version galGal5.

## Supplementary Materials II

**Supplementary Figure 1.** Local contact chromatin domains within GGA2_r1 region revealed by three different TAD calling algorithms (directionality index – black line, Armatus – green line, and TADTree – blue line) on Hi-C map of chicken embryonic fibroblasts (according to Fishman *et al.* 2019).

**Supplementary Figure 2.** FISH-mapping of DNA-probes to the regions of chicken lampbrush chromosomes GGA1_r1, GGA2_r1, GGA2_r2, GGA4_r1, GGA14_r1.

**Supplementary Figure 3.** FISH-mapping of DNA-probes to the loci within two neighboring interphase TADs in the region GGA4_r2 on chicken lampbrush chromosome 4 with double loop bridges.

**Supplementary Figure 4.** FISH-mapping of DNA-probes to the regions GGA4_r3 and GGA4_r4 on chicken lampbrush chromosome 4.

**Supplementary Figure 5.** FISH-mapping of DNA-probes to the regions GGA4_r3 and GGA4_r4 on highly decondensed lampbrush chromosome 4.

