## Supplementary Materials I for "Comparison of the somatic TADs and lampbrush chromomere-loop complexes in transcriptionally active prophase I oocytes"

**Supplementary Table 1.** Positions of BAC-clones within studied regions in the chicken genome assembly version galGal5

| Region name | BAC name | BAC start position | BAC end position | BAC total length, bp | Distance between 1st-2nd/2nd-3rd BACs, bp |
|---|---|---|---|---|---|
| GGA1_r1 | 178I7 | 146768336 | 146959667 | 191332 | 603519/586261 |
| | 191J12 | 147563186 | 147752805 | 189620 | |
| | 90G18 | 148339066 | 148550462 | 213397 | |
| GGA2_r1 | 177D19 | 25424085 | 25632182 | 208098 | 707043/697065 |
| | 96F4 | 26339255 | 26564682 | 225428 | |
| | 91I5 | 27261747 | 27445467 | 183721 | |
| GGA2_r2 | 179E9 | 45912588 | 46116014 | 203427 | 596080/590670 |
| | 182I12 | 46712094 | 46921822 | 209729 | |
| | 133G24 | 47512492 | 47718789 | 206298 | |
| GGA4_r1 | 177B7 | 5725062 | 5925483 | 200422 | 493658/497640 |
| | 33C6 | 6419141 | 6610711 | 191571 | |
| | 48P9 | 7108351 | 7324973 | 216623 | |
| GGA4_r2 | 49F17 | 37602285 | 37797874 | 195590 | 603228/602539 |
| | 96N10 | 38401102 | 38600730 | 199629 | |
| | 2P8 | 39203269 | 39396146 | 192878 | |
| GGA4_r3 | 108G8 | 40173462 | 40387074 | 213612 | 502057/467691 |
| | 34P18 | 40889131 | 41079977 | 190846 | |
| | 47F11 | 41547668 | 41747405 | 199737 | |
| GGA4_r4 | 14D4 | 62107450 | 62312871 | 205421 | 504204/515312 |
| | 139O9 | 62817075 | 62995554 | 178479 | |
| | 164E15 | 63510866 | 63683957 | 173091 | |
| GGA14_r1 | 31C6 | 9567659 | 9764723 | 197065 | 601235/614320 |
| | 22J21 | 10365958 | 10553851 | 187894 | |
| | 119E18 | 11168171 | 11366145 | 197975 | |

**Supplementary Table 2.** Summary of FISH-mapping data of BAC-clone DNA-probes to lampbrush chromatin domains with information on gene content according to chicken genome assembly version galGal5

| Lampbrush chromatin domains according to FISH mapping | Region name | BAC name | Number of annotated genes | Protein coding gene(s) | ncRNA gene(s) |
|---|---|---|---|---|---|
| Lateral loop | GGA2_r1 | 96F4 | 1 | TSHD7A | 0 |
| | GGA4_1 | 177B7 | 1 | DIAPH1 | 0 |
| | GGA4_2 | 49F17 | 1 | COL25A1 | 0 |
| | GGA4_3 | 108G8 | 4 | WWC2<br>DCTD | LOC107051751<br>LOC107051752 |
| | GGA4_3 | 34P18 | 0 | 0 | 0 |
| | GGA4_4 | 164E15 | 6 | MTMR7<br>VPS37A<br>CNOT7<br>ZDHHC2<br>MICU3<br>FGF20 | 0 |
| | GGA14_r1 | 119E18 | 1 | RBFOX1 | 0 |
| **Total** | **6** | **7** | **14** | **12** | **2** |
| Chromomere-loop complex | GGA2_r1 | 91I5 | 2 | ETV1 | LOC107052563 |
| | GGA2_r2 | 182I12 | 2 | AOAH<br>ANLN | 0 |
| | GGA2_r2 | 133G24 | 2 | NPSR1<br>FANCD2OS | 0 |
| | GGA4_2 | 96N10 | 2 | DKK2 | LOC107051761 |
| | GGA4_2 | 2P8 | 5 | CXXC4 | LOC107051758<br>LOC107051757<br>LOC107051756<br>miR1627 |
| | GGA4_3 | 47F11 | 0 | 0 | 0 |
| | GGA4_4 | 14D4 | 2 | FAT1 | LOC107053253 |
| | GGA14_r1 | 31C6 | 2 | 0 | LOC107054633<br>LOC101751275 |
| | GGA14_r1 | 22J21 | 0 | 0 | 0 |
| **Total** | **6** | **9** | **17** | **8** | **9** |
| Chromomere | GGA1_r1 | 178I7 | 1 | GPC6 | 0 |
| | GGA1_r1 | 191J12 | 1 | GPC5 | 0 |
| | GGA1_r1 | 90G18 | 0 | 0 | 0 |
| | GGA2_r1 | 177D19 | 0 | 0 | 0 |
| | GGA2_r2 | 179E9 | 2 | STAC<br>DCLK3 | 0 |
| | GGA4_r1 | 33C6 | 2 | 0 | LOC107052109<br>LOC101748814 |
| | GGA4_r4 | 139O9 | 0 | 0 | 0 |
| **Total** | **5** | **7** | **6** | **4** | **2** |
| Chromatin nodule | GGA4_r1 | 48P9 | 0 | 0 | 0 |
| **Total** | **1** | **1** | **0** | **0** | **0** |
| **Total (General)** | **8** | **24** | **37** | **24** | **13** |

**Supplementary Table 3.** Summary of FISH-mapping data of BAC-clone DNA-probes to lampbrush lateral loops obtained from previous studies (Krasikova et al. 2012; Zlotina et al. 2012) with information on gene content according to chicken genome assembly version galGal5

| Chromosome | BAC-clone name | Genetic marker or GenBank accession number | Gene name | Gene length, kb | Genetic marker position in galGal5 | | DNA/DNA+RNA FISH Figure # in Zlotina et al. 2012 | DNA/RNA FISH Figure # in Krasikova et al. 2012 |
|---|---|---|---|---|---|---|---|---|
| GGA1p | WAG43G6 | MCW0112 | PDE3A | 243.492 | 65026223 | 65025973 | Fig. 2a | |
| GGA3q | WAG13D11 | CZ566991 | ----------- | ----------- | 5944007 | 5944620 | Fig. 3a | Fig. 5b |
| GGA11p | WAG12F3 | LEI0110 | NUP93 | 70.776 | 2170218 | 2170442 | Fig. 4d | Fig. 5d |
| GGA11q | WAG52K20 | ADL0123 | KIFC3 | 11.867 | 525118 | 525391 | Fig. 4a | |
| GGA12q | WAG40H21 | CZ564147 | CENPP | 118.901 | 3442523 | 3443029 | Fig. 5a | |
| GGA14q | WAG32F10 | CZ562612 | SDK1 | 360.337 | 3695102 | 3695437 | Fig. 6a | |
