## Supplementary Materials II for "Comparison of the somatic TADs and lampbrush chromomere-loop complexes in transcriptionally active prophase I oocytes"

**Supplementary Figure 1**

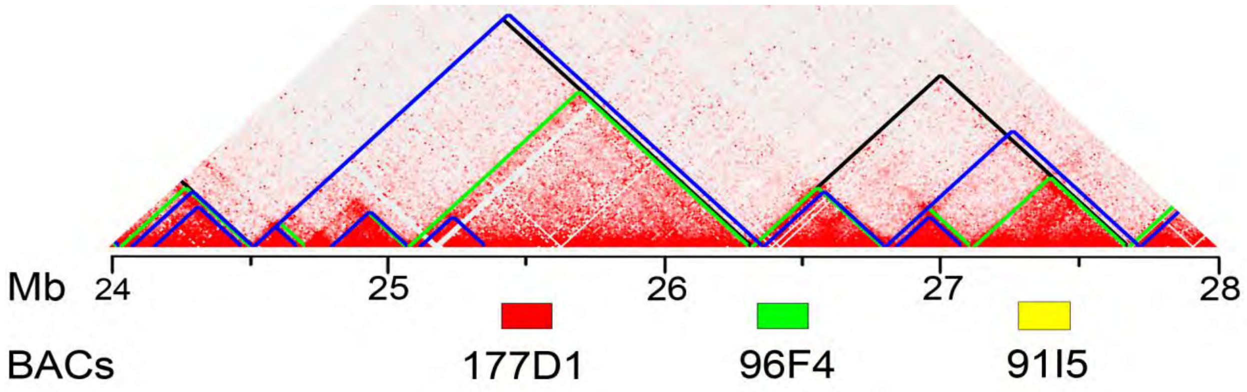


**Supplementary Figure 1.** Local contact chromatin domains within GGA2_r1 region revealed by three different TAD calling algorithms (directionality index – black line, Armatus – green line, and TADTree – blue line) on Hi-C heatmap of chicken embryonic fibroblasts (according to Fishman at al., 2019).

**Supplementary Figure 2**

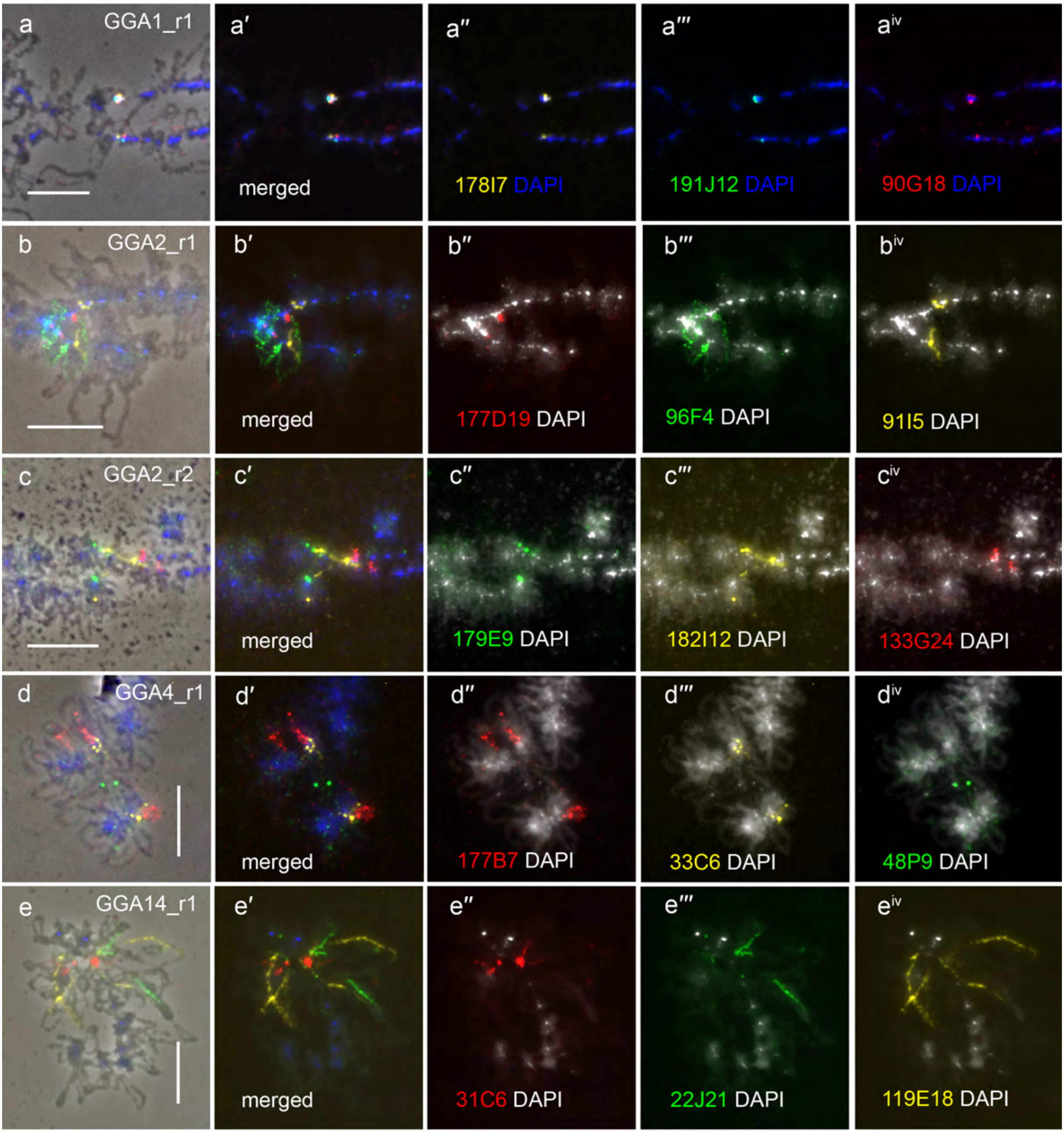

**Supplementary Figure 2. FISH-mapping of DNA-probes to the regions of chicken lampbrush chromosomes GGA1_r1, GGA2_r1, GGA2_r2, GGA4_r1, GGA14_r1.**

**a** - **a**$^{iv}$ – fragment of chicken lampbrush chromosome 1, FISH with DNA-probes to the loci of two neighboring TADs in the region GGA1_r1, **a** – fluorescent images merged with phase contrast image, **a′** – merged fluorescent images, **a″** - **a**$^{iv}$ – separate fluorescent images merged with DAPI, 178I7-Atto647N – yellow, 191J12-bio – green, 90G18 – red, DAPI – blue.

**b** - **b**$^{iv}$ – fragment of chicken lampbrush chromosome 2, FISH with DNA-probes to the loci of two neighboring TADs (three subTADs) in the region GGA2_r1, **b** – fluorescent images merged with phase contrast image, **b′** – merged fluorescent images, **b″** - **b**$^{iv}$ – separate fluorescent images merged with DAPI, 177D19-dig – red, 96F4-bio – green, 91I15-Atto647N – yellow, DAPI: (**b**, **b′**) – blue, (**b″** - **b**$^{iv}$) – white.

**c** - **c**$^{iv}$ – fragment of chicken lampbrush chromosome 2, FISH with DNA-probes to the loci of three neighboring TADs in the region GGA2_r2, **c** – fluorescent images merged with phase contrast image, **c′** – merged fluorescent images, **c″** - **c**$^{iv}$ – separate fluorescent images merged with DAPI, 179E9-bio – green, 182I12-Atto647N – yellow, 133G24-dig – red, DAPI: (**c**, **c′**) – blue, (**c″** - **c**$^{iv}$) – white.

**d** - **d**$^{iv}$ – fragment of chicken lampbrush chromosome 4, FISH with DNA-probes to the loci of two neighboring TADs in the region GGA4_r1, **d** – fluorescent images merged with phase contrast image, **d′** – merged fluorescent images, **d″** - **d**$^{iv}$ – separate fluorescent images merged with DAPI, 177B7-dig – red, 33C6-Atto647N – yellow, 48P9-bio – green, DAPI: (**d**, **d′**) – blue, (**d″** - **d**$^{iv}$) – white.

**e** - **e**$^{iv}$ – chicken lampbrush chromosome 14, FISH with DNA-probes to the loci of two neighboring TADs in the region GGA14_r1, **e** – fluorescent images merged with phase contrast image, **e′** – merged fluorescent images, **e″** - **e**$^{iv}$ – separate fluorescent images merged with DAPI, 31C6-dig – red, 22J21-bio – green, 119E18-Atto647N – yellow, DAPI: (**c**, **c′**) – blue, (**e″** - **e**$^{iv}$) – white. Scale bars – 10µm.

**Supplementary Figue 3**

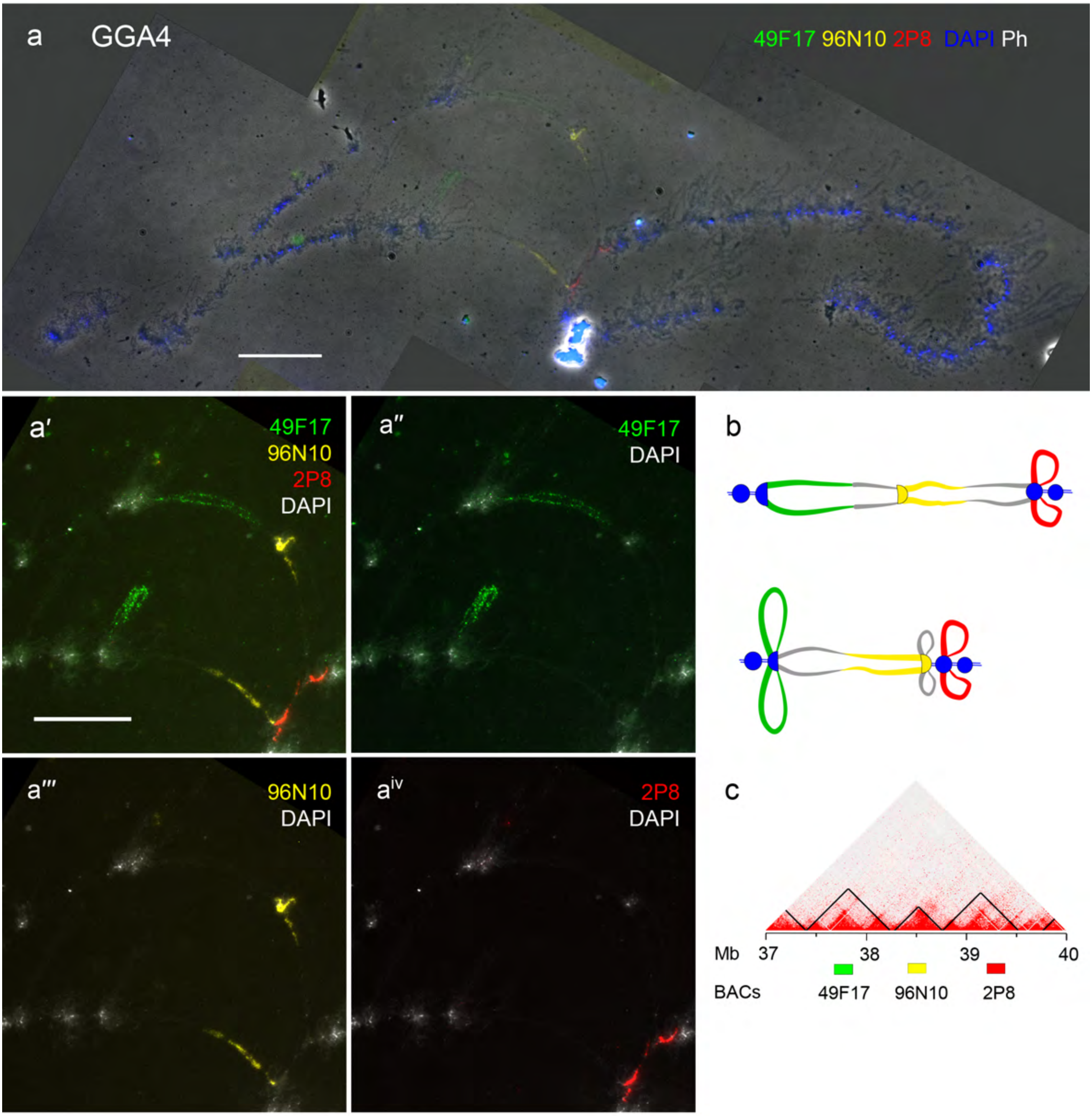

**Supplementary Figure 3. FISH-mapping of DNA-probes to the loci within two neighboring interphase TADs on chicken lampbrush chromosome region GGA4_r2 with double loop bridges. a - a$^{iv}$** – FISH with BAC-clone DNA-probes to the loci belonging to three neighboring interphase TADs in the region GGA4_r2 on chicken lampbrush chromosome 4 (GGA4); **a** – lampbrush chromosome 4 with the double loop bridges in the region GGA4_r2, fluorescent images merged with phase contrast image, 49F17-bio – green, 96N10-Atto647N – yellow, 2P8-dig – red, DAPI – blue, scale bar – 20 µm; **a′** - **a$^{iv}$** – enlarged images of the region GGA4_r2, DNA-probes colored as on **a** separately merged with DAPI (white), **a′** – merged fluorescent images, scale bar – 10 µm. **b** – schematic drawings of hybridization pattern with DNA-probes to interphase TADs loci in the region of GGA4_r2 on each homolog of the bivalent 4 presented on **a** - **a$^{iv}$**. **c** – Hi-C heatmap for chicken embryonic fibroblasts (CEF) for the region GGA4_r2 (according to Fishman et al., 2019), positions of BAC-clones indicated with boxes of the colors corresponding to that on panels **a** - **a$^{iv}$**.

**Supplementary Figure 4**

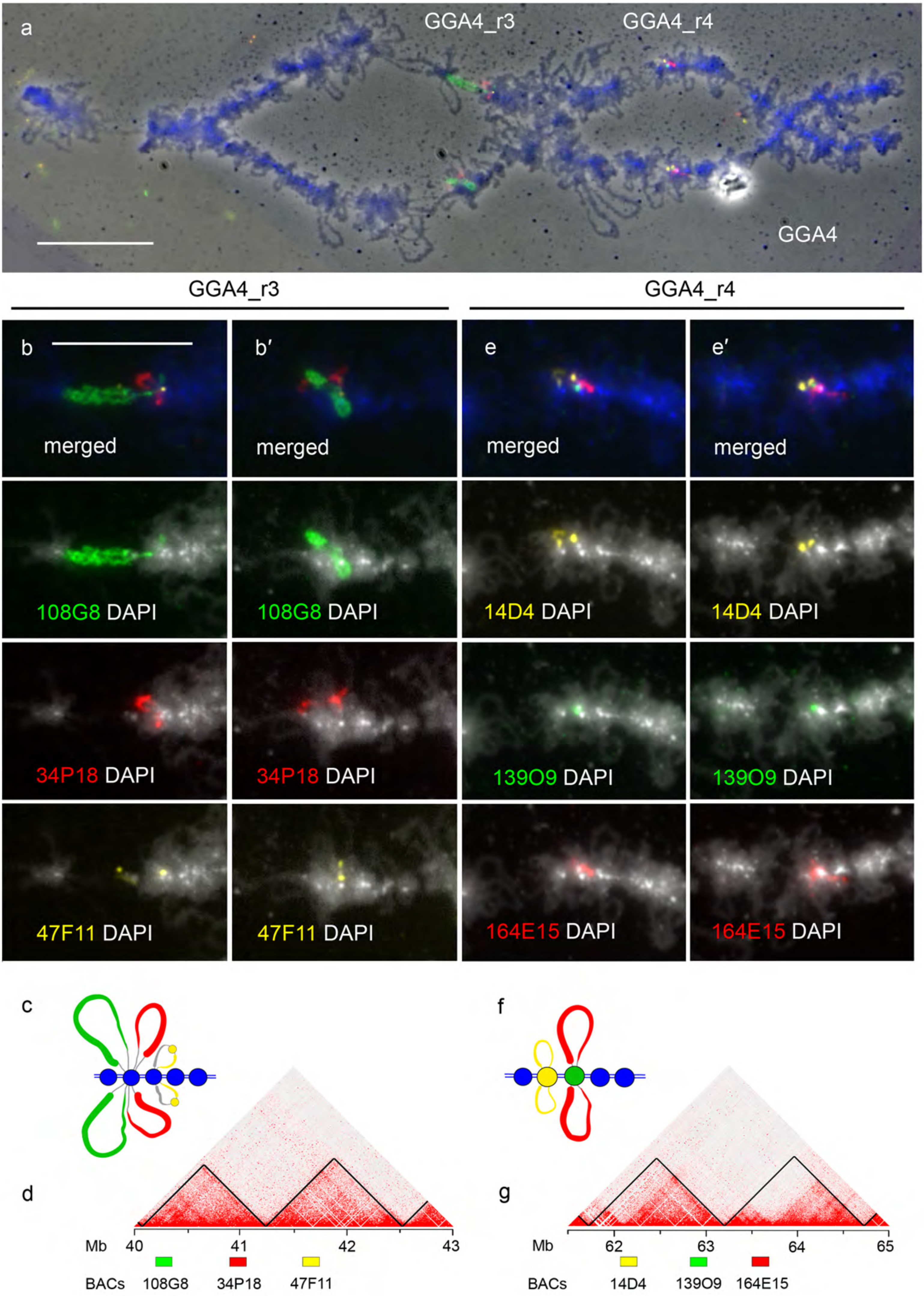

**Supplementary Figure 4. FISH-mapping of DNA-probes to the regions GGA4_r3 and GGA4_r4 on chicken lampbrush chromosome 4. a, b - b′, e - e′** – FISH with BAC-clone DNA-probes to the loci belonging to two pairs of neighboring interphase TADs in the regions GGA4_r3 and GGA4_r4. **a** – lampbrush chromosome 4, fluorescent image merged with phase contrast image, GGA4_r3: 108G8-bio – green, 34P18-dig – red, 47F11-Atto647N – yellow; GGA4_r4: 14D4-Atto647N – yellow, 139O9-bio – green, 164E15-dig – red, DAPI – blue, scale bar – 20 µm; **b - b′**, **e - e′** – enlarged fragments of the regions GGA4_r3 and GGA4_r4 correspondingly, DNA-probes colored as on **a**, top images of the panels – merged fluorescent images with DAPI colored blue, lower images – separate DNA-probe FISH merged with DAPI colored white, scale bar – 10 µm. **c**, **f** – schematic drawings of hybridization patterns with DNA-probes to interphase TAD loci on lampbrush chromosome 4 in the regions GGA4_r3 and GGA4_r4, correspondingly. **d**, **g** – Hi-C heatmaps for chicken embryonic fibroblasts (CEF) for the regions GGA4_r3 and GGA4_r4 (according to Fishman et al., 2019), positions of BAC-clones indicated with boxes of the colors corresponding to that on panels **a**, **b - b′**, **e - e′**.

**Supplementary Figure 5**

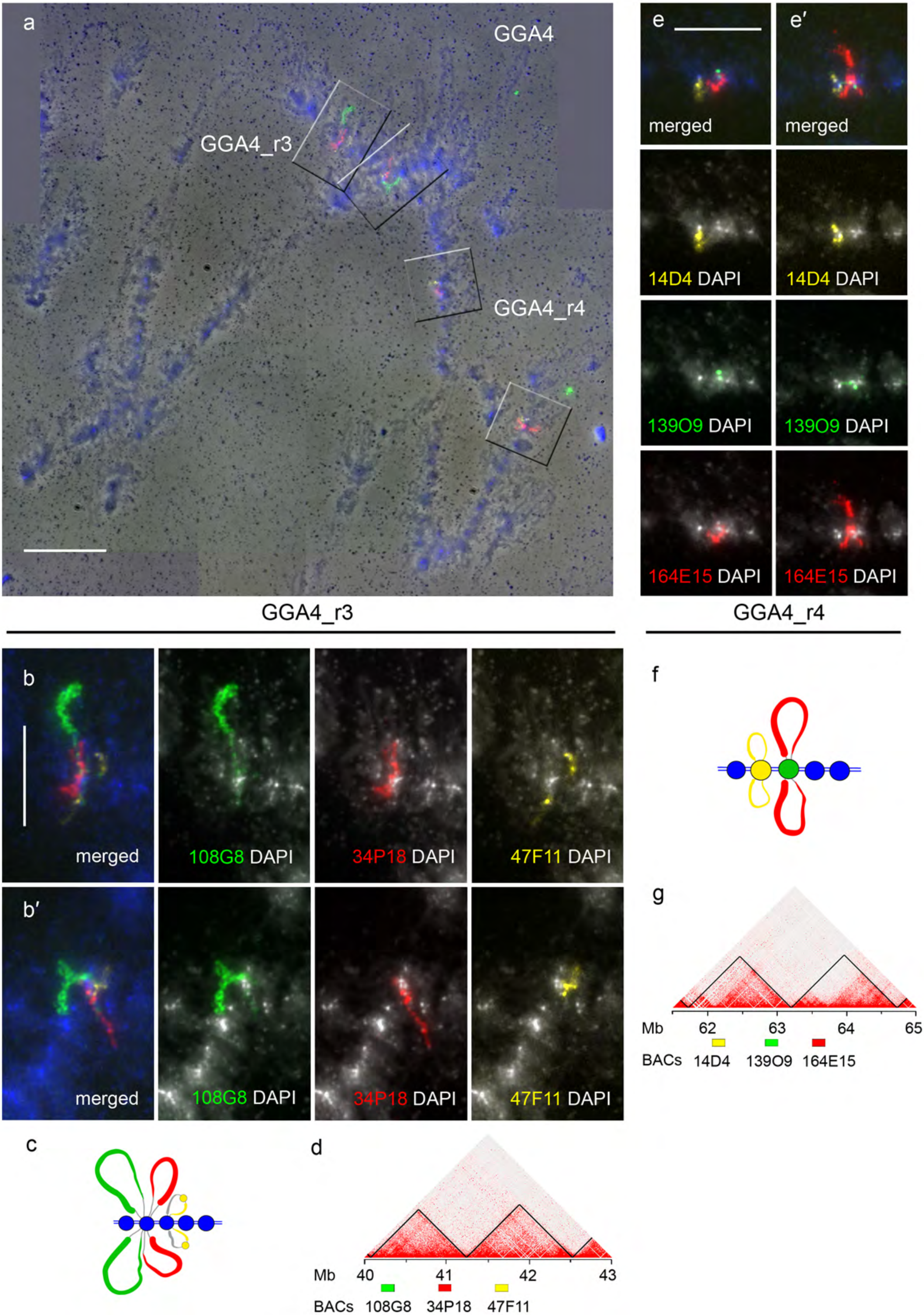

**Supplementary Figure 5. FISH-mapping of DNA-probes to the regions GGA4_r3 and GGA4_r4 on highly decondensed lampbrush chromosome 4.** DNA-probes to the loci belonging to two pairs of neighboring interphase TADs in the regions GGA4_r3 and GGA4_r4. **a** – lampbrush chromosome 4, fluorescent image merged with phase contrast image, GGA4_r3: 108G8-bio – green, 34P18-dig – red,47F11-Atto647N – yellow; GGA4_r4: 14D4-Atto647N – yellow, 139O9-bio – green, 164E15-dig – red, DAPI – blue, scale bar – 20 µm; **b** - **b′**, **e** - **e′** – enlarged fragments of the regions GGA4_r3 and GGA4_r4 correspondingly, DNA-probes colored as on **a**, top images of the panels – merged fluorescent images with DAPI colored blue, lower images – separate DNA-probe FISH merged with DAPI colored white, scale bar – 10 µm. **c**, **f** – schematic drawings of hybridization patterns with DNA-probes to interphase TAD loci on lampbrush chromosome 4 in the regions GGA4_r3 and GGA4_r4, correspondingly. **d**, **g** – Hi-C heatmaps for chicken embryonic fibroblasts (CEF) for the regions GGA4_r3 and GGA4_r4 (according to Fishman et al., 2019), positions of BAC-clones indicated with boxes of the colors corresponding to that on panels **a**, **b** - **b′**, **e** - **e′**.
